# Elevational Range Sizes of Woody plants Increase with Climate Variability in the Tropical Andes

**DOI:** 10.1101/2023.02.21.529430

**Authors:** Flavia Montaño-Centellas, Alfredo F. Fuentes, Leslie Cayola, Manuel J. Macía, Gabriel Arellano, M. Isabel Loza, Beatriz Nieto-Ariza, J. Sebastián Tello

**Affiliations:** Department of Biological Sciences, Louisiana State University, Baton Rouge, LA 70803, USA; Herbario Nacional de Bolivia, Instituto de Ecología, Carrera de Biología, Universidad Mayor de San Andrés, La Paz, Cajón Postal 10077, Bolivia; Missouri Botanical Garden, St. Louis, MO 63110, USA; Departamento de Biología (Botánica), Universidad Autónoma de Madrid, Calle Darwin 2, ES–28049 Madrid, Spain; Centro de Investigación en Biodiversidad y Cambio Global (CIBC-UAM), Universidad Autónoma de Madrid, Calle Darwin 2, ES–28049 Madrid, Spain; Department of Ecology and Evolutionary Biology, University of Michigan, Ann Arbor, MI 48109, USA; Oikobit LLC, Albuquerque, NM 87120, USA; Global Tree Conservation Program and the Center for Tree Science, The Morton Arboretum, Lisle, IL, 60532-1293, USA; Wetland Ecology, Dep. of Geography and Geoecology, Karlsruhe Institute of Technology, Karlsruhe, Germany; Center for Conservation and Sustainable Development, Missouri Botanical Garden, St. Louis, MO 63110, USA

**Keywords:** Andes, Bolivia, Climate variability hypothesis, Elevation, Forest plots, Madidi, Range size, Trees

## Abstract

**Aim:** The climate variability hypothesis proposes that species subjected to wide variation in climatic conditions will evolve wider niches, resulting in larger distributions. We test this hypothesis in tropical plants across a broad elevational gradient; specifically, we use a species-level approach to evaluate whether elevational range sizes are explained by the levels of thermal variability experienced by species.

**Location:** Central Andes

**Time period:** Present day

**Major taxa studied:** Woody plants

**Methods:** Combining data from 479 forest plots, we determined the elevational distributions of nearly 2300 species along an elevational gradient (∼209 – 3800 m). For each species, we calculated the maximum annual variation in temperature experienced across its elevational distribution. We used phylogenetic generalized least square models to evaluate the effect of thermal variability on range size. Our models included additional covariates that might affect range size: body size, local abundance, mean temperature and total precipitation. We also considered interactions between thermal variability and mean temperature or precipitation. To account for geometric constraints, we repeated our analyses with a standardized measure of range size, calculated by comparing observed range sizes with values obtained from a null model.

**Results:** Our results supported the main prediction of the climate variability hypothesis. Thermal variability had a strong positive effect on the range size, with species exposed to higher thermal variability having broader elevational distributions. Body size and local abundance also had positive, yet weak effects, on elevational range size. Furthermore, there was a strong positive interaction between thermal variability and mean annual temperature.

**Main conclusions:** Thermal variability had an overriding importance in driving elevational range sizes of woody plants in the Central Andes. Moreover, the relationship between thermal variability and range size might be even stronger in warmer regions, underlining the potential vulnerability of tropical montane floras to the effects of global warming.

## 1 INTRODUCTION

Assessing the mechanisms shaping the distribution of species is essential to better understand the assembly of local communities and the potential consequences of environmental drivers on biodiversity patterns (Bellard *et al*., 2012; Nadeau *et al*., 2017). This is particularly urgent for mountain regions, areas of great importance for biodiversity conservation (Jung *et al*., 2021). Mountain ranges are characterized by high environmental heterogeneity across space and time (Rahbek *et al*., 2019a), and harbor roughly one third of terrestrial living organisms, including many small-ranged species (Rahbek *et al*., 2019b). Understanding the drivers of montane species ranges, particularly in the tropics, is critical given the threats that climate change and human modifications of the landscape poses to the distribution and persistence of species in these regions.

Although species distributions are often modeled as a function of average environmental conditions in a site or a region, temporal variation in these conditions can have profound effects on population and species adaptations, and consequently on their distribution. The climate variability hypothesis proposes that species subjected to wider temporal variation in climatic conditions will evolve tolerances to broader environmental niches, resulting in wider geographic distributions (Stevens, 1989). Correspondingly, species experiencing stable environments would develop narrow tolerances and small geographic distributions. The climate variability hypothesis has been proposed as a potential explanation for some classical patterns in ecology and biogeography. For example, the increases in range size with latitude (Rapoport, 1982) could be a consequence of increases in seasonal or daily climatic variability toward temperate regions (Stevens, 1989; Chan *et al*., 2016). Similarly, Janzen (1967)’ classic proposition that tropical mountains represent physiologically stronger filters for organisms than temperate mountain could also be seen as a special case of the climate variability hypothesis. Janzen’s hypothesized that having evolved in less variable environments, montane tropical species will likely have limited acclimation responses and, in consequence, smaller elevational ranges than species in temperate mountains.

Within tropical mountains climatic variability can fluctuate significantly across elevation; for example, daily temperature variation can be dramatic at high elevations, but only mild in the lowlands (McCain, 2009). If temporal variation in climate influences species distributions within mountains, then species near tropical mountain tops should have more extensive elevational distributions than species in the lowlands. This extension of the climate variability hypothesis to elevation (ECVH; Stevens, 1992) has been tested in many taxa, producing conflicting results (McCain & Knight, 2013; Chan *et al*., 2016; Shah *et al*., 2021). Whereas some studies show increases in range size with latitude and elevation (e.g., Patterson *et al*., 1996; Pintor *et al*., 2015), others have refuted these patterns (e.g., (Hawkins & Felizola Diniz-Filho, 2006; Maccagni & Willi, 2022). Contradictory results have fueled a debate regarding whether species responses to climate variation is only a local phenomenon or a consistent pattern (Rohde, 1996; McCain & Knight, 2013). Part of the reason for the inconsistent results among studies testing ECVH could be limitations to analyses or data. Geometric constraints in the distribution of species, for instance, have rarely been accounted for when testing ECVH hypothesis, resulting on strong criticisms on the statistical approaches and assumptions behind these studies (Ribas & Schoereder, 2006; McCain, 2009; Macek *et al*., 2021). Additionally, most studies of ECVH carry out assembly-level analyses, where the average range size for groups of species is used, and analyses focus on how these averages change across space (Rohde, 1992; Stevens, 1992). However, the evolution of climatic tolerances and responses to climatic variability are species-specific and, as such, require species-level analysis. Species-level and high-quality datasets are rare, particularly in diverse tropical regions, preventing species-level tests of ECVH in tropical mountains.

Studies testing predictions from the ECVH in vascular plants include only a few examples on the sub-tropical floras of the Himalayas (e.g., Liang *et al*., 2021; Macek *et al*., 2021) and the temperate Caucasus mountains (Mumladze *et al*., 2017), as well as studies focusing on smaller sets of temperate plants that extend ECVH predictions to other species characteristics (e.g., trait plasticity; Molina-Montenegro & Naya, 2012; Maccagni & Willi, 2022). To our knowledge the only study that has directly tested the climate variability hypothesis on Neotropical plants has focused only on alpine communities in the Andes (>3000 m; Cuesta *et al*., 2020). This study found that tropical alpine species had narrower thermal niches than temperate species, supporting predictions derived from the classic CVH, but no significant effects of elevation. In consequence, whether temporal climatic variability shapes the distribution of tropical plant species along elevational gradients is yet poorly understood. A thorough analysis of the predictions of ECVH is necessary, particularly in tropical plants, to understand how species respond to climatic variability, and particularly, whether climatic variability can promote the formation of large geographic ranges.

Here, we present the most thorough species-level test of the climate variability hypothesis across elevations (ECVH). Specifically, we evaluate whether species with populations experiencing high levels of temperature variation will have larger elevational distributions. For our study, we use data from the Madidi Project (www.madidiproject.weebly.com), a collaborative effort to document and study the plant biodiversity of the Tropical Andes, and possibly the most extensive dataset on tree occurrences in any tropical mountain. Using this data, we evaluate the effect of climate variability while controlling for the effects of geometric constraints and the potential confounding effects of other factors that have been proposed to affect range size, such as species characteristics (e.g., size), species abundance, and local temperature and precipitation. Our dataset and approach provide a unique opportunity to study the forces that drive the distribution of tree species in one of the most species-rich regions of the planet.

## 2 METHODS

### 2.1 Vegetation Data and Elevational Range Sizes

Species elevational distributions were determined based on a large network of forest plots distributed along the eastern slopes of the Bolivian Andes (Fig. 1). The network consists of 48 1-ha plots (henceforth large plots) and 458 0.1-ha plots (small plots) ranging in elevation from 209 m (Amazon forests) to 4,347 m (tree line). Within plots, all individuals of woody plant species with diameters at breast height (dbh) equal or above 10 cm (for large plots) or 2.5 cm (for small plots) were measured and identified. Each individual tree was assigned a species or morphospecies name (henceforth simply species), and extensive taxonomic work was conducted to ensure that names were applied consistently throughout all plots. For this study, we used version 5.0 of the Madidi Project plot database (https://doi.org/10.5281/zenodo.5160379). The species-level data and code necessary to replicate our analyses has been deposited and can be freely accessed in Zenodo ([*link will be updated upon submission*]).

**Figure 1.**
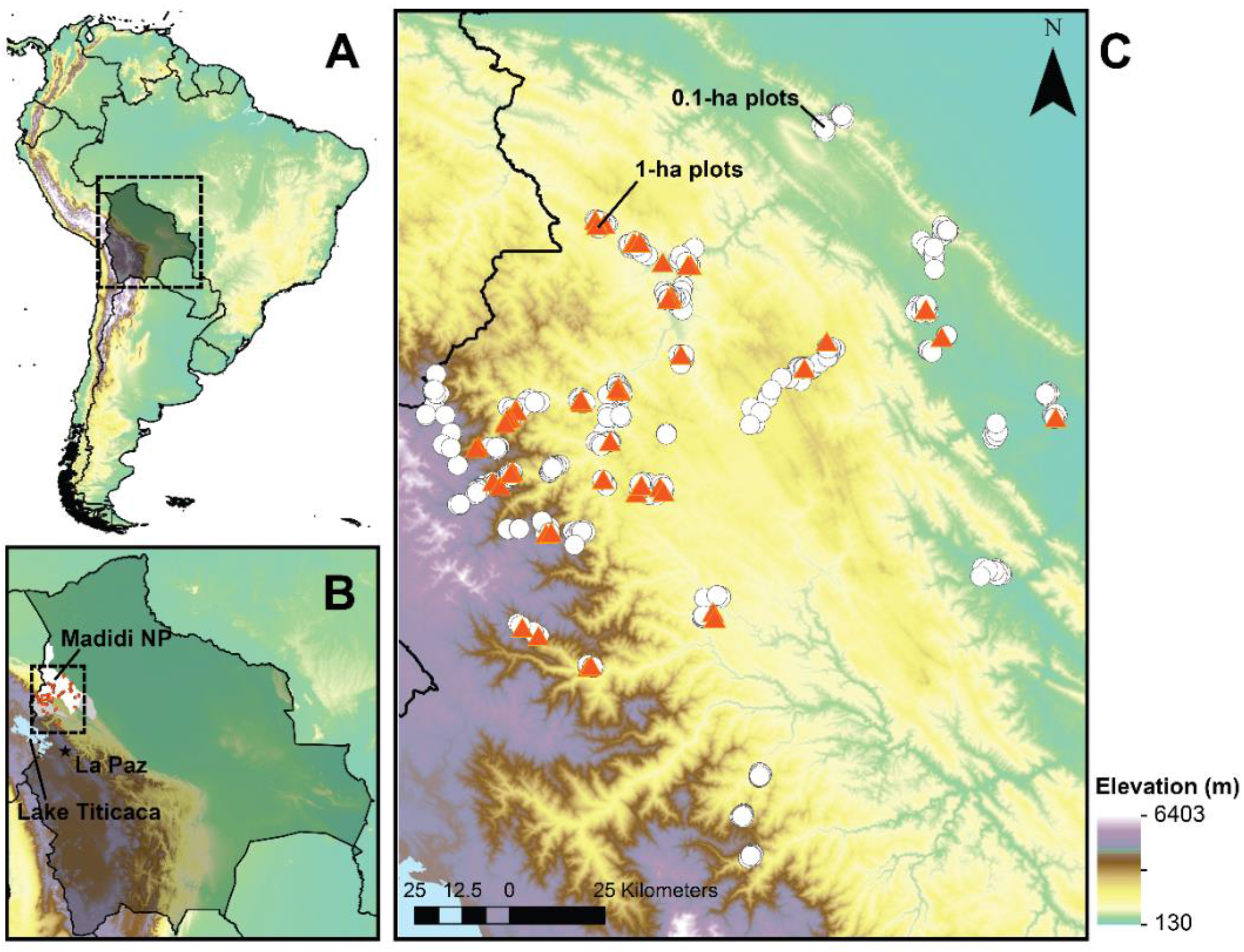
Map of the study region and network of forest plots. **(A & B)** Location of the study region within and around Madidi National Park in Bolivia. **(C)** The forests ‘ plots dataset used in our analyses include 48 large plots (1-ha in area) and 458 small plots (0.1-ha).

From these data, we removed all cacti (Cactaceae); bamboos (Poaceae), tree ferns (Dicksoniaceae and Cyatheaceae), gymnosperms (Podocarpaceae), and the non-native genera *Eucalyptus* and *Coffea*. We also removed plots above 3,800 m in elevation, which were dominated by species of *Polylepis* and likely managed by local communities. Finally, because we only sampled individuals with a dbh 2.5 cm or larger, species that rarely reach this size might be present in our data but seriously under-represented relative to their true abundances. Thus, we examined the distribution of species-level maximum size values across our dataset and eliminated all species with maximum size below the lowest 5% of the distribution (this is, all species with maximum size less or equal to 3.24 cm; see Fig. S1). This resulted in the elimination of 169 individuals of 126 species. Finally, we eliminated 1,328 individuals that could not be assigned to species or morphospecies (<1% of individuals) and 436 individuals from 16 additional species that could not be placed in the regional phylogeny (see below). After data curation, our dataset contained information on the distribution of 153,084 individuals belonging to 2,292 species across 479 forest plots (48 large plots and 431 small plots).

We estimated the elevational distribution of each species in our dataset by recording their highest and lowest elevation of occurrence (Fig. 2A). Elevational range size was calculated as the difference in meters between these two points. Elevational position was characterized by the species’ elevational mid-point (the mean between the maximum and minimum elevation of occurrence). For the estimation of elevational distributions, we merged the data from large and small plots with the objective of using all the available information for each species. After this process, we further excluded 461 species that were found in only one plot and thus could not be assigned to a reliable value of elevational range. Elevational range size had a strongly skewed frequency distribution (Fig. 2B). Most species had very small ranges and very few had broad elevational distributions. The mean and median of elevational range size were 702 and 607 m respectively; the maximum was 2,812 m.

**Figure 2.**
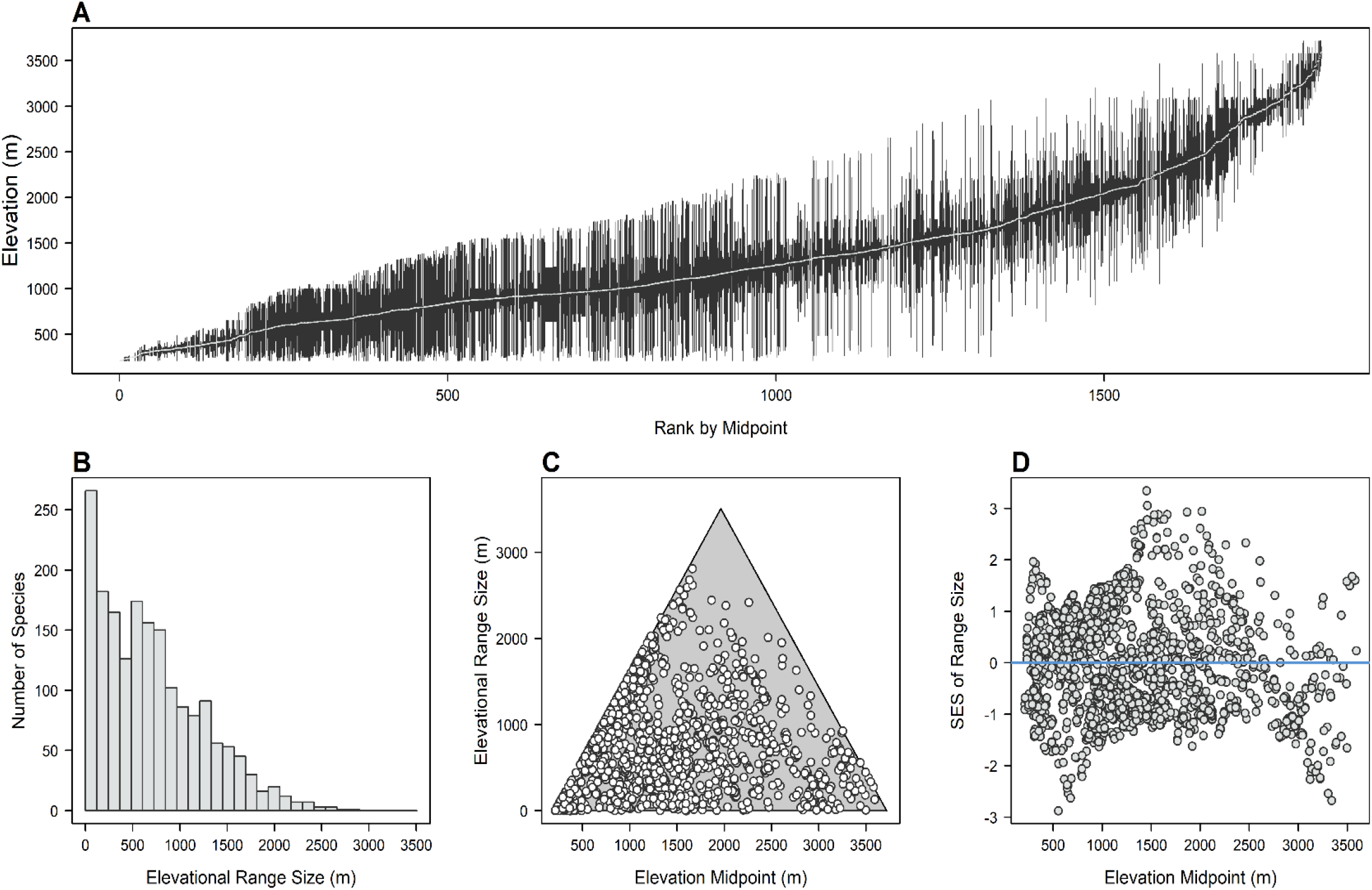
Elevational ranges for woody plant species in the Andes of northwestern Bolivia. **(A)** Vertical lines connect the lowest and highest elevations recorded for each species used in our analyses. In this way, the lines describe the elevational extent of species ‘ distributions (i.e., their elevational ranges). Species are ranked in the x-axis by their elevational mid-point of distribution. **(B)** Frequency distribution of elevational range sizes showing that most species have small ranges. **(C)** Relationship between elevational range size and elevational mid-point. The range size of each species is constrained geometrically by its position with respect to the upper and lower limits of the gradient. Species that have an elevational mid-point in the lowlands or highlands are constrained to having small ranges, while species that have a mid-point at intermediate elevations are free to have either small or large range sizes. The grey area shows the possible distribution of points; the black edges mark the geometric limits to this relationship. **(D)** Relationship between the standardized effect size of range size and elevational mid-point. Standardized effect size for a species was calculated by comparing its empirical range size to a subset of ranges of similar geometric constraints (see Methods for details & Figure 3). SES values greater than zero indicate ranges that are larger than expected by their constraints.

To account for heterogeneity among species in range size estimation and ensure that our conclusions did not depend on the precise collection of species used, we repeated all analyses using two alternative subsets of species (Fig. S2). In the second set of species, we included only species that were present in 3 or more sites or that had 5 or more individuals (1,713 species); in the third set, we subsampled forest plots to reduce heterogeneity in sampling effort across elevations. We divided the elevational gradient (209 to 3,717 m) into 20 equal-sized bands; in each band, we randomly selected 12 forest plots. This procedure reduced the data to 220 forest plots and 71,165 individuals, resulting in 1,280 species included in the third dataset. Despite considerable differences among these datasets, all analyses support the same conclusions, thus we present results for our complete dataset in the main text and provide results for the two subsets of data in the supplementary materials.

### 2.2 Temperature and Precipitation Data and Species-Level Predictors

Using the coordinates of each forest plot, we extracted temperature data from WorldClim 2.0 (at ∼1km resolution; Fick & Hijmans, 2017). We considered other alternative climate datasets (e.g., Chelsa 1.2; Karger *et al*., 2017), but we found that WorldClim 2 was the closest match to field measurements of temperature (Fig. S3). We extracted precipitation data from TRMM 2b31-Based Rainfall Climatology Version 1.0 at ∼1km resolution; Mulligan, 2006). For each plot, we obtained data on mean annual temperature (MAT), total annual precipitation (TAP), annual temperature range (ATR) and diurnal temperature range (DTR). Across the plot network, mean annual temperature decreases dramatically across elevations from 25.4 to 9 °C (Fig. S4A). Similarly, estimates of total precipitation range from 3,819 to 197 mm per year. Although temperature variability generally increased with elevation (Fig. S4B), the pattern was non-linear: annual temperature range showed a small initial dip towards intermediate elevations (with a minimum around 1,250 m), before a steep increase towards the highlands. Finally, diurnal temperature range is closely correlated with annual temperature range (Fig. S4D). For this reason, the effect of annual and diurnal variability could not be disentangled. All analyses use only data on annual temperature range, but similar models were produced when diurnal temperature range was used instead (Table S1). These gradients in climate suggest that populations of species in the highlands experience a higher degree of temperature fluctuations than in the lowlands. The distribution of plots across environmental gradients is depicted in Fig. S5 and S6.

For each species, we estimated the degree of temperature variation that individuals experience by using the maximum value of annual temperature range at a site across all occupied plots (Max. ATR). Additionally, we calculated other species-specific predictors that could be important determinants of elevational range size, which were used as co-variables in our analyses. We calculated abundance-weighted mean annual temperature (w-MAT) and total annual precipitation (w-TAP). For these calculations, plot-level values of MAT or TAP contribute to the species mean as a function of the abundance of the species in each plot. These variables represent the most typical environmental conditions occupied by each species. Finally, we calculated species-level maximum size as the 90 % quantile of the distribution of diameters at breast height (DBH) for each species, and species abundance as the maximum value of relative abundances of each species across all occupied plots.

### 2.3 Statistical test of hypotheses

The climate variability hypothesis across elevations (ECVH) predicts that species with populations experiencing high levels of temperature variation will have larger elevational distributions. To evaluate this prediction, while accounting for shared evolutionary history among species, we used a phylogenetic generalized least squares (PGLS) regression model, where species elevational range size was the dependent variable and maximum annual temperature range (max. ATR) was the main predictor of interest. In this analysis, errors were modeled using a Pagel correlation structure, which is more flexible than a Brownian correlation. Phylogenetic relationships among our species are based on Smith and Brown ‘s (2018) mega-phylogeny, accessed using the R package V.PhyloMaker (Jin & Qian, 2019). Species that were not found in the base phylogeny were added using taxonomic information at base of the branch of the corresponding genus or family using the “S1” option in V.PhyloMaker. While this phylogeny is a coarse description of evolutionary relationships, it allows us to construct phylogenetic regressions that would otherwise be impossible. We used Ives’ proposed R2resid metric to characterize the amount of variance in the data explained within a phylogenetic regression model (Ives, 2019); rr2 R package: (Ives & Li, 2018). PGLS models were performed with function gls in R package nlme (Pinheiro *et al*., 2020).

To account for the effects of other potentially important covariates, the PGLS model also included maximum size (i.e., 90th percentile of dbh distribution per species), species abundance, mean annual temperature (w-MAT) and total annual precipitation (w-TAP). To meet model assumptions, elevational range size was square-root transformed, while maximum size and species abundance were log-transformed. Other variables remained untransformed. All predictors were centered to a mean of zero and standardized to a standard deviation to 1 before analyses. In this way, regression coefficients are comparable and measure the relative importance of each predictor in the model. Finally, the model also included the interactions of temperature variability with mean temperature (max. ATR × w-MAT) and annual precipitation (max. ATR × w-TAP). We evaluated collinearity among predictors in our PGLS model using variance inflation factors (VIF) using function vif in the R package car (Fox *et al*., 2022). Most variables had VIFs less than 5 indicating that collinearity is low in our models (Table 1).

**Table 1.** Testing for the effect of temperature variability and other predictors on size of elevational distributions. Phylogenetic generalized least-square regressions (PGLS) were used. Elevational range size (ERS) or a standardized effect size for range size (SES) were modeled as the response variable in separate models. Regardless of the response used, we found that maximum annual temperature range (max. ATR) was a strong predictor and had a significant interaction with abundance-weighted mean annual temperature (w-MAT). Additional predictors included species maximum size, maximum abundance among occupied plots, and abundance-weighted total annual precipitation (w-TAP). For each predictor, we report standardized coefficients, p-values and variation inflation factors (VIF). Model fit is characterized by Ives ‘ residual R2 value for phylogenetic modes, as well as Pearson ‘s correlation between observed and model-predicted values of the response variable. Finally, we used a likelihood ratio test (LRT) to obtain a model-wide p-value by contrasting each PGLS against a null model. The null model had only an intercept and the same phylogenetic structure estimated for the main PGLS model. Details on univariate models and other competing models are detailed in Table S1.

| Response | Predictor | Coeff. | P-value | VIF | Obs.-Pred.<br>Corr. | $R^2_{\text{resid}}$ | LR | LRT:<br>P-value |
| --- | --- | --- | --- | --- | --- | --- | --- | --- |
| ERS | Intercept | 27.011 | < 0.001 |  |  |  |  |  |
|  | log(Max. Size) | 1.688 | < 0.001 | 1.054 |  |  |  |  |
|  | log(Max. Abund.) | 3.033 | < 0.001 | 1.156 |  |  |  |  |
|  | w-MAT | 1.730 | < 0.001 | 2.069 |  |  |  |  |
|  | w-TAP | 2.602 | < 0.001 | 1.952 | 0.618 | 0.389 | 890.6 | < 0.001 |
|  | Max. ATR | 9.592 | < 0.001 | 4.071 |  |  |  |  |
| | w-MAT × Max. ATR | 3.735 | < 0.001 | 5.877 | | | | |
| | w-TAP × Max. ATR | 0.309 | 0.310 | 3.591 | | | | |
| SES | Intercept | 0.237 | < 0.001 |  |  |  |  |  |
|  | log(Max. Size) | 0.136 | < 0.001 | 1.055 |  |  |  |  |
|  | log(Max. Abund.) | 0.252 | < 0.001 | 1.157 |  |  |  |  |
|  | w-MAT | 0.474 | < 0.001 | 2.035 |  |  |  |  |
|  | w-TAP | 0.257 | < 0.001 | 1.946 | 0.647 | 0.419 | 994.5 | < 0.001 |
|  | Max. ATR | 0.968 | < 0.001 | 4.014 |  |  |  |  |
| | w-MAT × Max. ATR | 0.280 | < 0.001 | 5.806 | | | | |
| | w-TAP × Max. ATR | 0.018 | 0.494 | 3.550 | | | | |

### 2.4 Accounting for Geometric Constraints and Sampling Effects on Elevational Range Sizes

Geographic ranges are subject to geometric constraints given by the limits of the domain over which species are distributed. In our study, the domain is the elevational gradient ranging from Amazon forests at 209 m in elevation and the timberline at 3,717 m. Species in our study are constrained to be distributed between these elevations (Fig. 2C). The effect of this constrained domain is expressed in the relationship between elevational position and elevational range size. Species with distributions centered in the lowlands (low elevational mid-points) or in the highlands (high elevational mid-points) cannot have large elevational ranges. Species with distributions centered at intermediate elevations, on the other hand, are free to have small or large elevational distributions. This constraint is potentially problematic for our analysis and could mask the effects of environmental or biological variables on the extent of the geographic distribution of species.

To account for this potential effect, we calculated an alternative metric of range size that is less affected by these geometric constraints. First, for a focal species, we calculated the distance d between its elevational mid-point to the closest edge of the elevational domain (i.e., to 209 m or 3,717 m whichever is closest; Fig. 3A). This distance determines the strength of the geometric constraint on a species’ distribution; as this distance decreases, the range of possible elevational range sizes decreases (Fig. 3A). Then, we found a pool of other species under similar geometric constraints. This pool was defined as all species with d values equal to that of the focal species ± 50 m (Fig. 3A and B). Species close to both ends of the domain of distribution can contribute to this pool of species with similar constraints (Fig. 3A). We calculated a standardized effect size (SES) that compares the elevational range size of the focal species to those of the other species in its pool (Fig. 3B). SES is simply the difference between the elevational range size of the focal species and the mean range size of all other species in its pool divided by the standard deviation of the range sizes in the pool. A positive SES value indicates that the focal species has a distribution that is larger than other species under similar constraints, while a negative value means the species has a smaller distribution. SES values were calculated in this way for all species. The statistical analyses described above for elevational range size were repeated using SES of range size.

**Figure 3.**
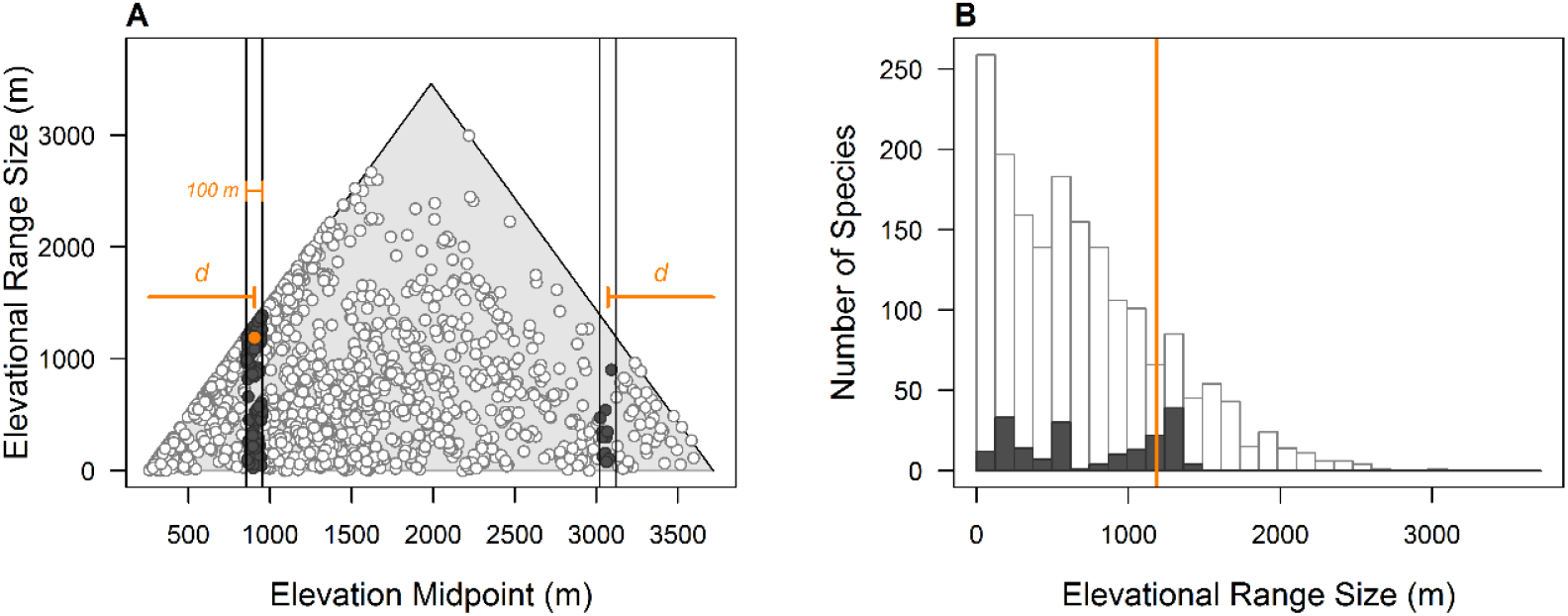
Calculation of standardized effect sizes of range size. To account for geometric constraints on elevational distributions, we compared the range size of each species to the range sizes of all other species that suffer from a similar constraint. In the example shown here, the range size and midpoint of a focal species is indicated by the orange circle in **(A)** and the vertical orange line in **(B)**. All species that suffer a similar constraint to the focal species are indicated by dark gray circles in **(A)** and gray bars in **(B)**. All other species are shown in white. Species with a similar constraint to the focal species are those that (1) have a midpoint in a region 50 m above or below the midpoint of the focal species, or (2) have a midpoint 50 m above or below an elevation that is equidistant from the opposite edge of the gradient (distance *d*). The range sizes of all species in these elevational bands represent a pool of potential values that the focal species could take given its midpoint. Thus, to calculate a standardized effect size, we (1) sampled 1,000 values of range size from the pool of similar species, and then (2) subtracted the mean of the random values from the empirical range size and divided this by the standard deviation of the random distribution. In this way, a standardized effect size measures the breadth of elevational distribution while accounting for geometric constraints. A positive value indicates a range size that is larger than other ranges with similar constraints; a negative value indicates a range size that is smaller than other similar ranges.

Finally, it is possible that relationship between range size and climate variability could be spuriously produced by a sampling effect; species with large elevational ranges might also occupy many sites (high occupancy). In turn, species present in many sites are able to sample the environmental space better and - by chance - find higher values of climatic variability (e.g., max. ATR). To account for this potential effect, we (1) examined the relationships between species occupancy (number of plots with presence of the species) and elevational range size and max. ATR, and (2) repeated our PGLS regressions including occupancy as a covariate. We found no evidence of this potential sampling bias in our analyses; while high occupancy does lead to larger ranges (Fig. S7A), high occupancy does not necessarily imply larger values of temperature variability (Fig. S7B). Moreover, the main conclusions of our analyses did not vary when including occupancy as a covariate in our PGLS models (Table S2).

## 3 RESULTS

Our results provide strong evidence that elevational range sizes are shaped by temporal variation in climate, particularly temperature. We found that maximum annual temperature range was the strongest predictor included in our models (Table 1) and had a clear positive effect on elevational range size (Fig. 4). Species exposed to higher temperature variability have broader geographic distributions. This effect was highly consistent across our different analyses; maximum annual temperature range had a strong positive effect whether elevational range size or standardized effect sizes were used as response variables (Fig. 4; Fig. S8). Similarly, maximum annual temperature range had a consistent positive effect when elevational distributions were characterized using species with at least 2 occurrences (Fig. 4, Table S1), or when using species with at least 3 occurrences or 5 individuals (Fig. S9, Table S3), or with a dataset that has been reduced to homogenize effort across elevations (Fig. S10, Table S4).

**Figure 4.**
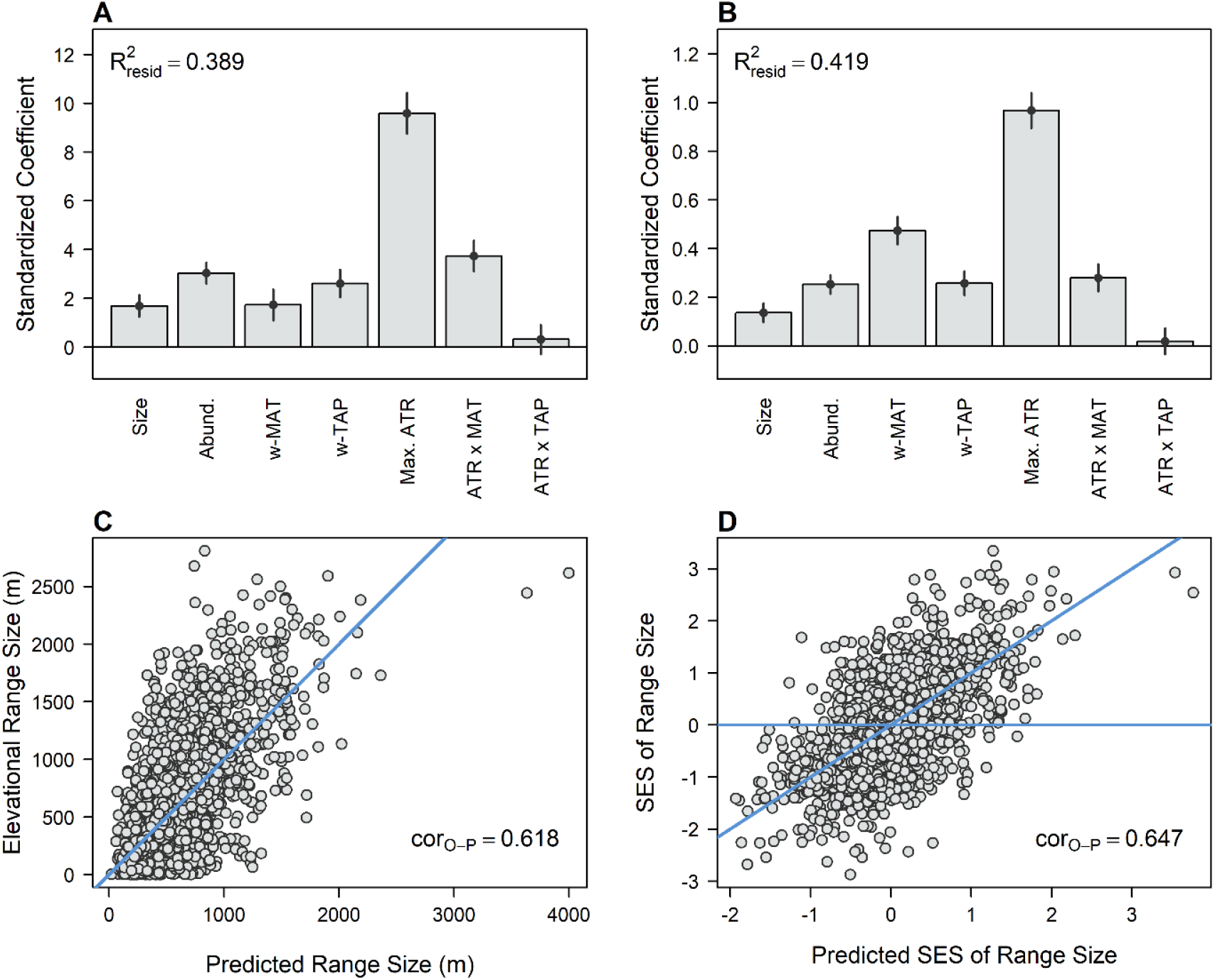
Effects of climate variability and other predictors on the breadth of elevational distributions. **(A)** Standardized coefficients showing the effect of each predictor on elevational range size. The height of each bar indicates the coefficient estimate, while the lines show the 95% confidence interval. Ives ‘ R^2^ is also shown. **(B)** Same as (A), but where the response variable was the standardized effect sizes of range size (i.e., range size after accounting for geometric constraints). **(C)** Empirical values of range size plotted against predictions made by the regression model in (A). The 1:1 correspondence is indicated by the solid blue line. **(D)** Same as (C), but where the response variable was the standardized effect sizes of range size (regression model in B). Size: maximum size (90^th^ percentile of diameter at breast height); Abund.: maximum local relative abundance; w-MAT: abundance-weighted mean annual temperature; w-TAP: abundance-weighted total annual precipitation; ATR: maximum annual temperature range.

We found evidence that temperature variability interacts with mean annual temperature, but not with total annual precipitation (Fig. 4 A and B). While the effect of temperature variability is always positive, the strength of this effect is greater for species with distributions in warmer climates (Fig. 5A and B). On the other hand, the effect of annual temperature range is consistent regardless of levels of precipitation (Fig. 5C and D). Finally, several other species characteristics had a significant effect on elevational range size, but the effect sizes were small (Fig. 4; Fig. S8). Range size increased for larger species (maximum size), and species that were more locally common (maximum abundance). Furthermore, species had larger elevational ranges in warmer and more humid places. These results were also robust when using alternative datasets.

**Figure 5.**
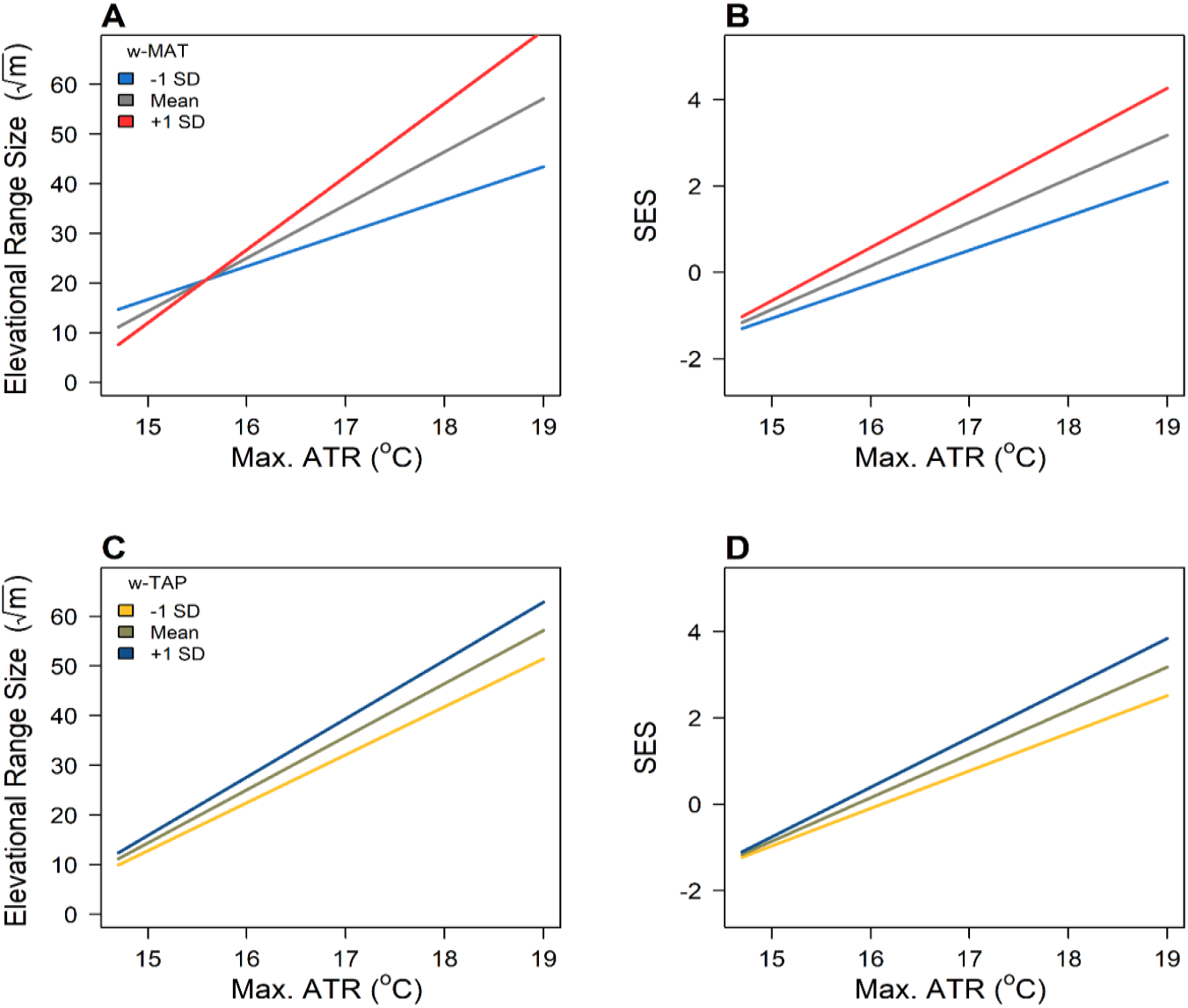
Interactions between temperature variability and mean temperature or total precipitation. Each panel shows the effect of temperature variability (maximum annual temperature range; max. ATR) on elevational range size (left column) and standardized effect sizes (right column) for different values of mean annual temperature (top row) and total annual precipitation (bottom row). In each case, the gray line depicts the effect of max. ATR for the mean value of the interacting variable. The colored lines depict the effects of max. ATR for values one standard deviation above and below the mean of the interacting variable. These results demonstrate that increases in mean temperature significantly amplify the effect of temperature variability **(A & B)**. On the other hand, increases in total precipitation do not modify the effects of temporal variability **(C & D)**.

## 4 DISCUSSION

### 4.1 Thermal variability and mean temperature interact to determine elevational range size

Using species-specific responses for ∼2300 plant species to climate variability across an extensive elevational gradient in the Central Tropical Andes, we found strong support for the climate variability hypothesis across elevations (ECVH; (Janzen, 1967; Stevens, 1992). Our findings show a strong positive relationship between local climate variability, particularly in temperature, and the elevational range size of woody plants (Table 1). In fact, the effect of variability in temperature is stronger than that of any other factor considered in our models. Importantly, our results were robust to all variations in analyses to account for potential biases related with species rarity and range size variability, uneven sampling across elevations and geometric constraints. The overall trend for elevational restricted species to occupy less climatically variable environments, regardless of their elevation of occurrence (i.e., not limited to lower elevations), suggests that elevational range restriction in Andean trees is likely related to narrow thermal tolerances rather than to biotic interactions or habitat specificity, two processes hypothesized to be more prevalent at lower elevations (MacArthur, 1984; Brown *et al*., 1996; Paquette & Hargreaves, 2021).

Previous studies have tested for the relationship between thermal variability and elevational range size, independent of the elevation of occurrence. Like ours, these studies found this relationship to be posited, despite using different methods of assessing climatic variability and focusing on different taxa (Pintor *et al*., 2015; Beck *et al*., 2016; Maccagni & Willi, 2022). Only a few studies have, however, tested predictions from the ECVH in vascular plants and their conclusions have been limited by their data or analyses. For instance, using a large empirical dataset on plant elevational distributions in the Western Himalaya, (Macek *et al*., 2021) found no support for the ECVH. As recognized by the authors, the lack of relationship between climate variability and elevational ranges in their study might result from the fact that the lowest elevation in their study is ∼2650 m a.s.l, and thus their dataset lacks information of lowland species and lower elevation climatic variability. A similar reason might have caused the lack of relationship between thermal niche breadth (maximum – minimum temperature a species experienced) and elevation in alpine plants (> 3000 m a.s.l.) of the Andes in the study by Cuesta et al. (2019). Here, we take advantage of a naturally extreme elevational gradient (∼ 200 - ∼3750 m a.s.l.) in the Central Tropical Andes and can extend our hypothesis testing to the whole set of woody plants. To our knowledge, no other comparable dataset exists for tropical plants where sampling of species has been as intensive and systematic over a large elevational gradient, and empirical data was obtained with standardized and homogeneous taxonomic information across species and sites. When including a full gradient of climatic variability, we found a strong positive relationship between thermal variability and the elevational range sizes.

While we found that climate variability has a strong positive effect on elevational range size, we also found that the magnitude of this effect depends on whether species are distributed in warmer or colder regions (i.e., a significant interaction between maximum annual temperature range and abundance-weighted mean annual temperature). Specifically, the positive effect of climate variability was stronger for species in warmer regions (e.g., lower elevations) than for species in colder regions (e.g., higher elevations). This finding is consistent with previous studies that found that both mean climatic conditions and climate variability are important drivers of species’ range sizes in different taxa (e.g., (Luo *et al*., 2011; Chan *et al*., 2016; Liang *et al*., 2021). Although studies in terrestrial vertebrates have considered the interaction between average environmental conditions and climate variability (Chan *et al*., 2016), to our knowledge our study is the first one on woody plants to include these effects when testing the ECVH. For example, Liang et al. (2021) considered mean environmental variables besides thermal variability in their study of plant elevational ranges. They found that both mean annual temperature and mean annual precipitation had a significant relationship with plant elevational ranges; they did not, however, consider interactions among these and climate variability in their analyses. Similarly, Mumladze et al. (2017), examined the correlations between the thermal range size of plant species (and not directly its elevational range) and the maximum temperature seasonality in two elevational gradients of the Caucasus. In their paper, Mumladze et al. (2017) separately tested the correlation with precipitation ranges and precipitation seasonality but did not test for interactions nor examined the relative importance of different environmental variables on species ranges. Our study, therefore, is the first one to show how environmental conditions modulate the effect of climate variability on the climatic tolerances and range size of plant species.

Although we argue that the most direct way to test predictions of the ECVH is to study the response of individual species to different levels of climate variability, most studies use the average range size of co-occurring species as response variable (e.g., Mumladze et al. 2017; Liang et al. 2021; Macek et al. 2021). These studies average the range sizes of all species occurring at a given site or elevational band (i.e., the “Steven’s method”) or average the range size of species whose distributional middle point falls within a given elevational band (i.e., the “midpoint method”). Because species relationships with climate variables is idiosyncratic (McCain & Knight, 2013), this aggregation of species responses could be a confounding factor, resulting in inconsistent results. Assemblage-level averages hide important variation among species. In a study with mayflies, for example Gill *et al*., (2016) found great variability in elevation range sizes even among closely related species that was likely related with variation in species physiological and dispersal traits. Species-specific differences in traits may result in large variation in elevational ranges across plant clades that co-occur at any given elevation, variability that could be dismissed when using assemblage-level metrics. Thus, conclusions reached with assemblage-level analyses should be taken cautiously.

### 4.2 Limitations of our study and recommendations for future analyses

A potential caveat of our study is the use of climate information from global databases. The coarse resolution of global databases might result in inaccurate information in mountain regions (e.g., (Browoski & Schickhoff, 2017). The complex landscape of the Andes likely adds to thermal variability; adjacent areas with different topographic exposures may differ notoriously in temperature and thermal variability, creating contrasting micro-habitat variation (Jackson & Forster, 2010). This small-scale spatial variability might be better captured with local climatic information obtained, for example, from data-loggers installed across elevations. We partially address this concern by comparing the climatic patterns in global datasets with a few data-loggers located in the study region (Fig. S3). We found WorldClim v.2 matched most closely the climate patterns we detected *in situ* with data-loggers. Furthermore, the use of highly localized climate data collected by data-loggers has its own limitations. With a forest canopy that might surpass 30 m of height, environmental information obtained from sensors located below 3 m (where most data loggers are set due to logistical constraints; (Bach *et al*., 2003) might represent poorly the thermal environment that most trees experience. Indeed, it has been shown that climatic conditions experienced by understory vs. canopy species can vary substantially (Frey *et al*., 2016), with canopies potentially experiencing greater temperature variability (De Frenne *et al*., 2019). Future studies might explore the differences in climate variability experienced by understory and canopy species and how these further affects species ‘ distribution across environmental gradients.

Finally, it is important to consider that although we found range sizes to be strongly associated with climate variability, other mechanisms might also act as determinants of Andean plant species’ ranges. We found a significant effect of tree size and local abundance, both of which had a positive effect on range size. These results are consistent with other studies on the ecological factors shaping the size of species distributions (Stahl *et al*., 2014). Moreover, other processes that we did not consider in our analyses could also be important. Biotic interactions such as specialized mutualisms or competitive interactions have been found to shape species ranges across latitudes and elevation (Brooker *et al*., 2007; Jankowski *et al*., 2010; Wisz *et al*., 2013). Dispersal abilities might also play a role in the realized range size of plants; with climate stability potentially having a stronger effect on groups with lower dispersal capacities (Xu *et al*., 2018). Finally, phenotypic plasticity or local adaptation can both contribute to shaping elevational range sizes(Bradshaw, 1965)(Van Nuland *et al*., 2017; Buckley *et al*., 2019). Future studies should focus on disentangling which and how these mechanisms might further restrict or extend species’ elevation distributions in tropical mountains.

### 4.3 Implications for environmental change

Understanding how climate shapes species distributions along environmental gradients is becoming increasingly urgent in a rapidly changing world, particularly because many tropical species are responding to global warming through range shifts (Nadeau *et al*., 2017; Fadrique *et al*., 2018; Freeman *et al*., 2018). Our study points to the overriding importance of thermal variability in driving elevational range sizes in woody plants in the Central Tropical Andes. Moreover, we found evidence suggesting that in warmer mountains the relationship between thermal variability and range size might be even stronger. Our results have implications not only to understand drivers of range size, but to predict how environmental change might impact biodiversity (Nadeau *et al*., 2017).

Combined, our findings highlight the great vulnerability of tropical floras to the enhanced effects of climate change in mountain ranges (Sentinella *et al*., 2020). Under a climate warming scenario, species with smaller thermal tolerances (often in warm, tropical regions) might be more vulnerable as their distributions seem to be strongly related with their climatic stability. Furthermore, species at low elevations not only have narrower climatic tolerances and small ranges, but they often also experience temperatures closer to their upper tolerance limits (Colwell *et al*., 2008). This could mean that species inhabiting tropical lowlands will likely face greater impacts of warming temperatures. Species might respond to such changes in local conditions either by tracking suitable climates and moving upslope to match their historical niches (e.g., Feeley *et al*., 2011), by persisting in situ in thermally buffered micro-habitats created by topography and other physiographic features (e.g., Suggitt *et al*., 2018), potentially resulting in fragmented populations, or by decreasing in abundance and potentially going extinct. By the same logic, species that are adapted to more variable environments and have broader distributions might be best able to cope with significant environmental change. On the other hand, as thermal variability along elevation is mostly determined by variation in minimum rather than maximum temperatures (lower temperatures decrease at a faster rate), species adapted to highly variable climates may struggle surviving in habitats with temperatures closer to their maximum tolerances. Having no place to “escape” from higher temperatures, high elevation floras in the tropics might be particularly vulnerable to rising temperatures. Climate variability, its effect on species climate tolerance and spatial distribution, can provide important clues into how species, communities and ecosystems will change in response to environmental shifts.

