## Additional Tables and Figures for "Elevational Range Sizes of Woody plants Increase with Climate Variability in the Tropical Andes"

### ADDITIONAL FIGURES

**Figure S1. Species with small maximum diameters that were removed from the data.** The figure shows the relationship between total numbers of individuals against the 90$^{th}$ percentiles of diameter at breast height (dbh; "maximum size") for species in our dataset. In our forest plots, we recorded individuals only with diameters (dbh) greater or equal to 2.5 cm in small plots and 10 cm in large plots (horizontal dashed lines). Species that infrequently reach 2.5 cm in diameter might appear rare despite potentially being common at smaller sizes, and their distributions might be poorly estimated from our data. To avoid this potential issue, we eliminated all species whose 90$^{th}$ percentile of dbh was less than 3.24 cm (red line and points). This limit of 3.24 cm was found as the 5$^{th}$ percentile of the distribution of 90$^{th}$ percentiles of dbh across species.

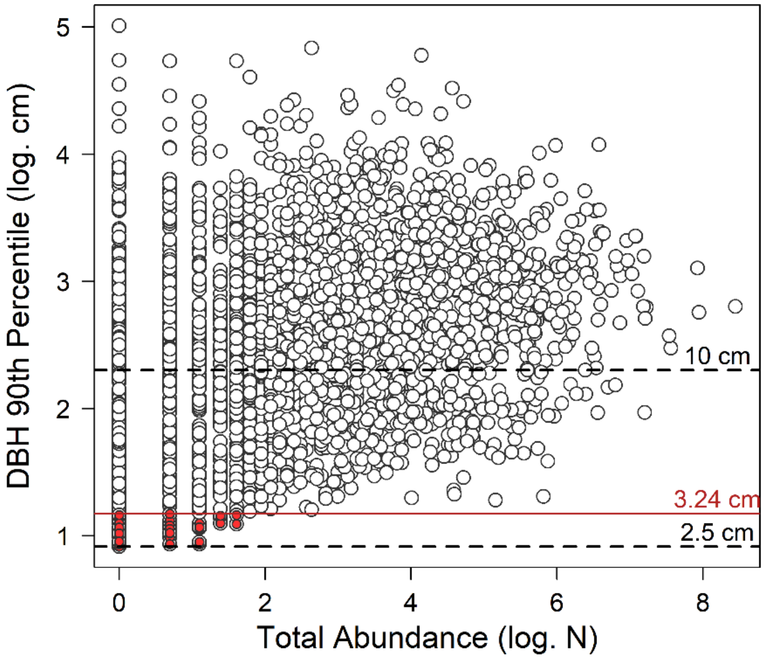

**Figure S2. Elevational range sizes of species in alternative datasets.** For different versions of the dataset, rows show the relationship between elevational range size and elevational midpoint (first column), as well as the frequency distribution of range sizes (second column). The first row (A&B) uses all species, including all those present at a single plot and those removed owing to other criteria (see Methods). The second row (C & D) shows the subset of species used for analyses in the main body of the manuscript. These species are present in at least 2 plots. The third row (E & F) shows a subset of species present in at least 3 plots or having at least 5 individuals. These first three rows use data from all plots (Figure 1 & S5). The last row (G & H), however, contains species present in at least 2 plots, but where the plot data has been modified to sample the elevational gradient homogeneously (Figure S6). Alternative analyses were conducted for data in third and fourth rows to check that our conclusions did not depend on how the dataset was constructed (Figures S7 & S8). These alternative analyses led to similar conclusions.

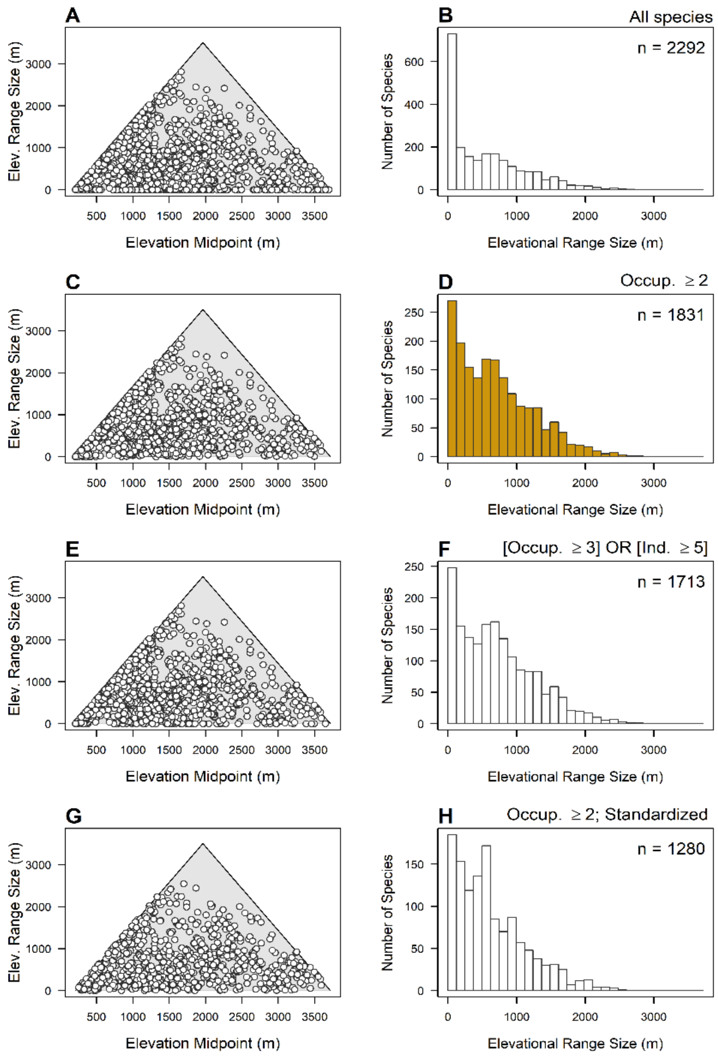

**Figure S3. Elevational gradients in temperature per plot across different climatic datasets.** The climatic dataset considered is represented by the grey dots, while the orange symbols correspond to field observations by HOBO U23 Pro v2 data loggers at 5 sites. The data logger observations range only from September 2011 to March 2012, so we used this range as a "year" in the calculations below. We also downloaded daily values of land surface temperature (LST) as measured by MODIS in the Terra satellite. For the LST dataset, we used information from January 1$^{st}$ 2002 to December 31$^{st}$ 2017. For each dataset, we obtained or calculated values of mean annual temperature, diurnal and annual temperature rage, as well as minimum and maximum temperature values. We found that WorldClim version 2.1 and LST data from MODIS-Terra have values that coincide well with field observations of annual temperature range. CHELSA 1.2 differed more significantly, so it was not considered further.

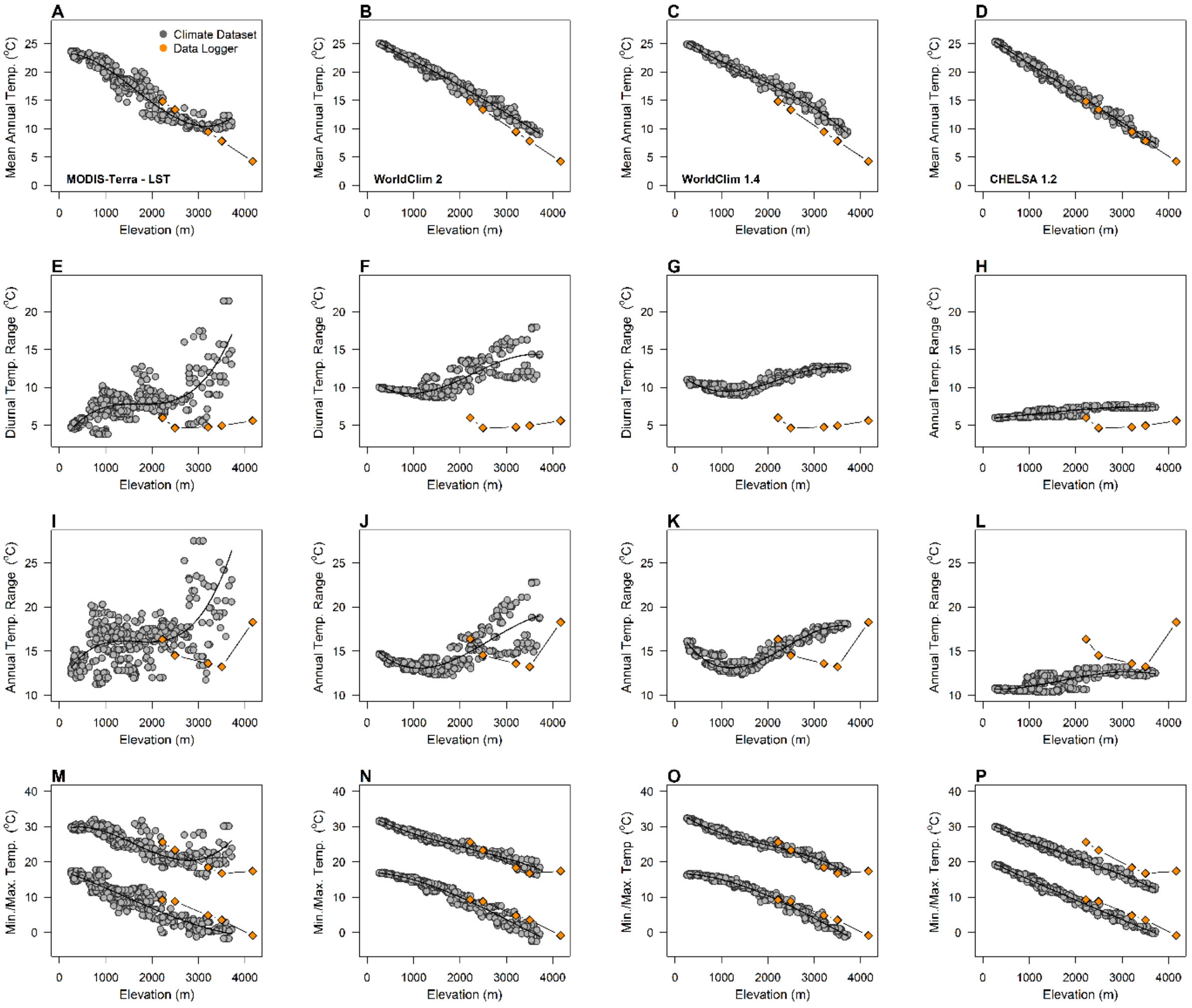

**Figure S4. Elevational gradients in temperature and temperature variability.** Patterns are presented separately for small plots (grey circles and solid line) and large plots (white circles and dashed line). **(A)** Mean annual temperature decreases while **(B)** annual temperature range increases with elevation across our plots in northwestern Bolivia. **(C)** Maximum and minimum temperatures at each site (i.e., maximum of the warmest month and minimum of the coldest month) both decrease with elevation, but lower temperatures decrease at a faster rate leading to an increase in annual temperature range towards higher elevations. **(D)** This annual range in temperature correlates closely with diurnal temperature range among our study plots. In **(C)**, vertical lines connect the maximum and minimum temperatures at a particular plot; the horizontal broken line marks the line of 0 °C. For this figure and our analyses, we used temperature data from Worldclim 2.1 (Fick & Hijmans, 2017, 2).

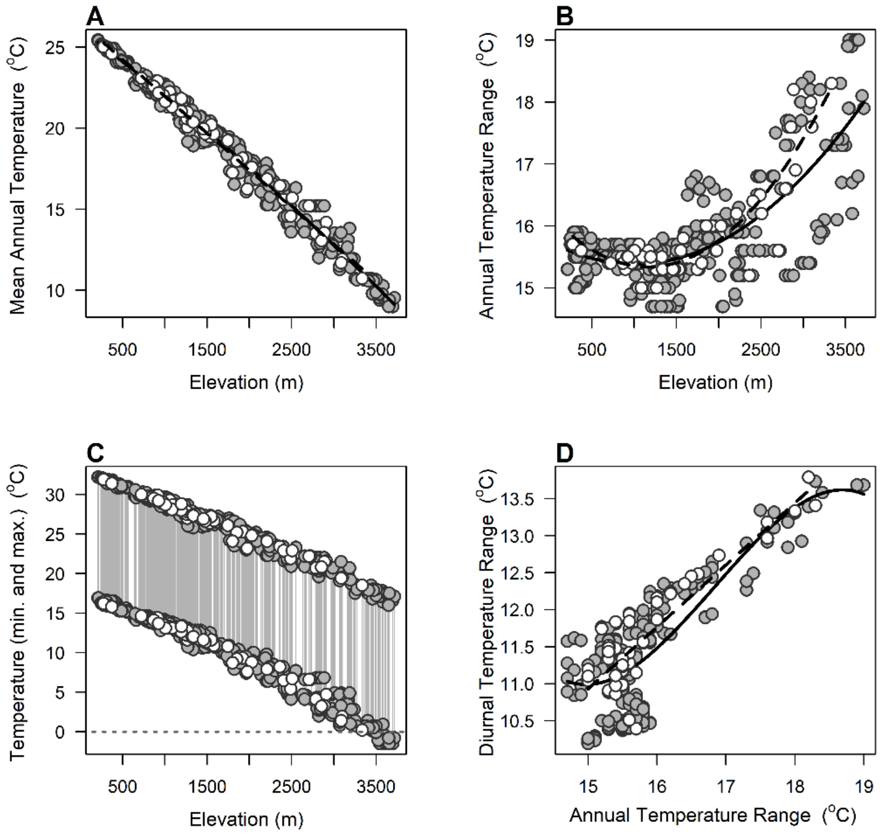

**Figure S5. Environmental distribution of forest plots.** The histograms show the frequency distribution of elevation **(A)**, mean annual temperature (MAT; **B**), total annual precipitation (TAP; **C**) and annual temperature range (ATR; **D**) across all forest plots in our dataset (Figure 1). These plots represent the sites used to estimate the abundance of each species, as well as their elevational and environmental distributions.

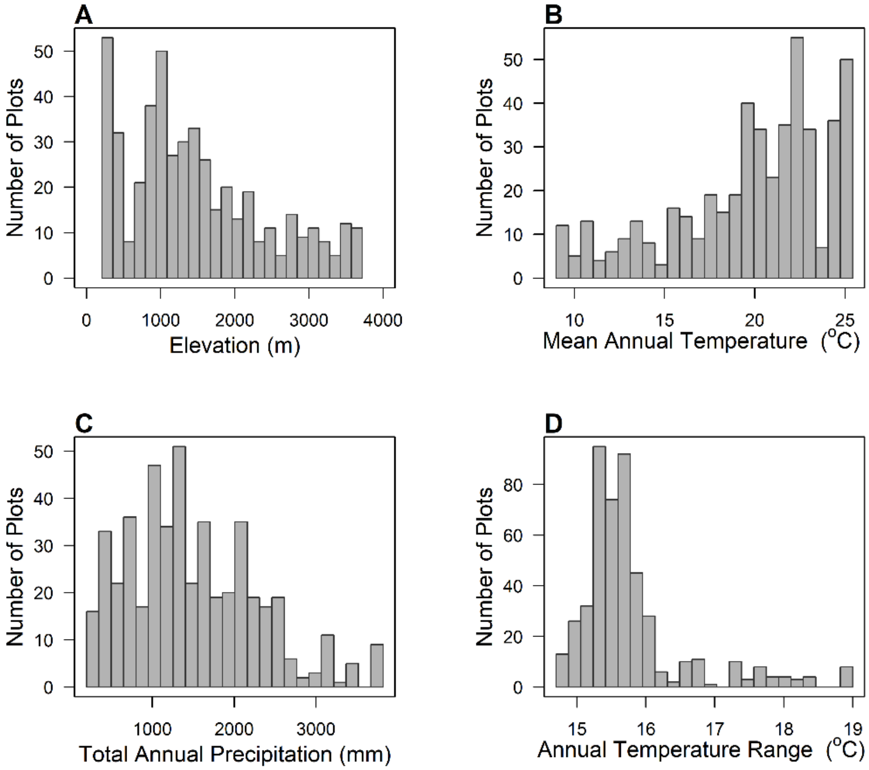

**Figure S6. Environmental distribution of forest plots when sampling across elevations has been standardized.** In this subset of the data, we divided the elevational domain (209 to 3,717m) into 20 equal-sized bands of 184.6 m. We then sampled 12 plots in each band or retained all plots when less than 12 were available. This prevented strong elevational biases in the distribution of plots (see Figure S6). The histograms show the frequency distribution of elevation **(A)**, mean annual temperature (MAT; **B**), total annual precipitation (TAP; **C**) and annual temperature range (ATR; **D**) across all forest plots in our dataset (Figure 1). These plots were used to estimate the abundance of each species, as well as their elevational and environmental distributions.

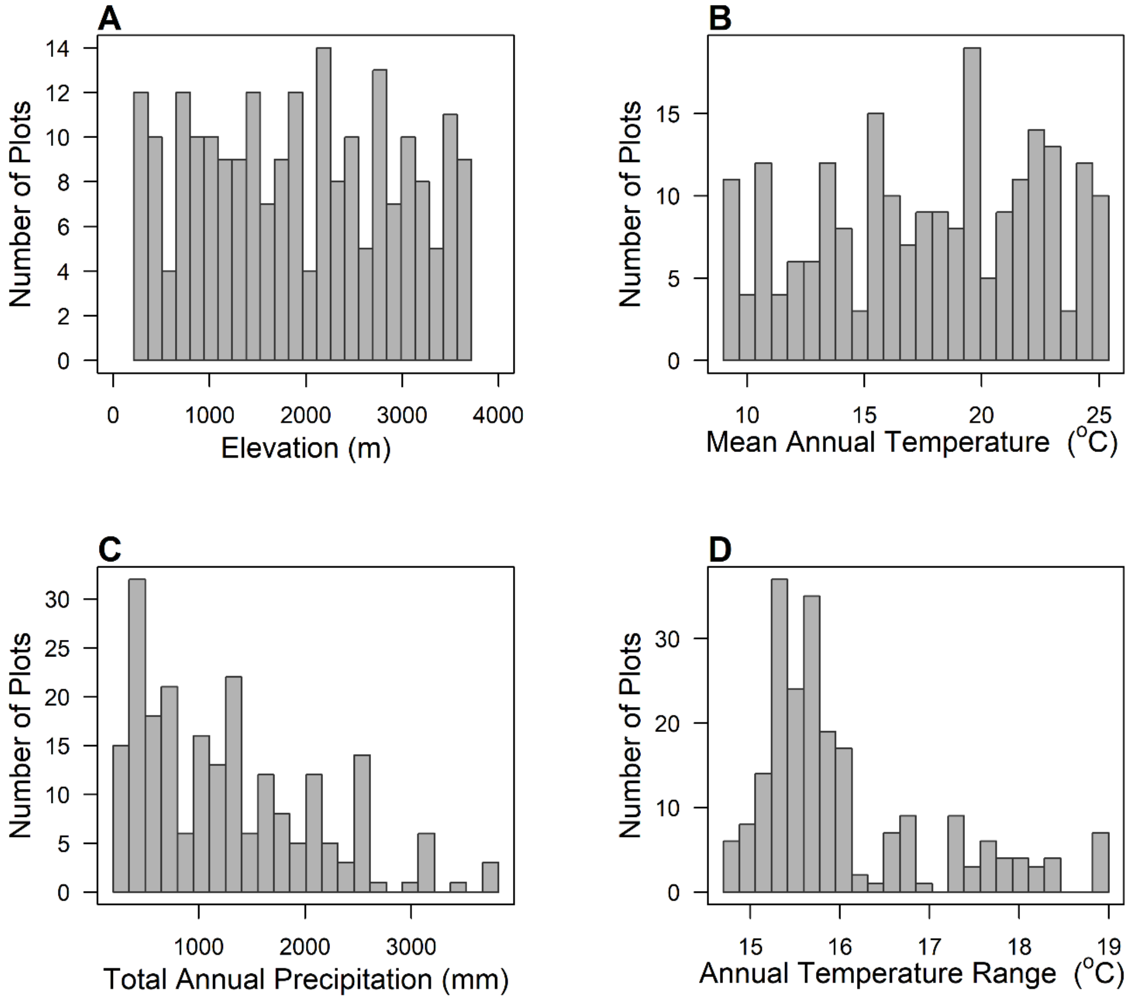

**Figure S7. Relationships between range size and climate variability with occupancy.** The relationship between range size and climate variability could be spuriously produced by a sampling effect. While high occupancy leads to larger ranges (A), high occupancy does not necessarily imply larger values of temperature variability (B). This potentially confounding effect of occupancy is also absent from regression models that include occupancy as one of the covariables (Table S2).

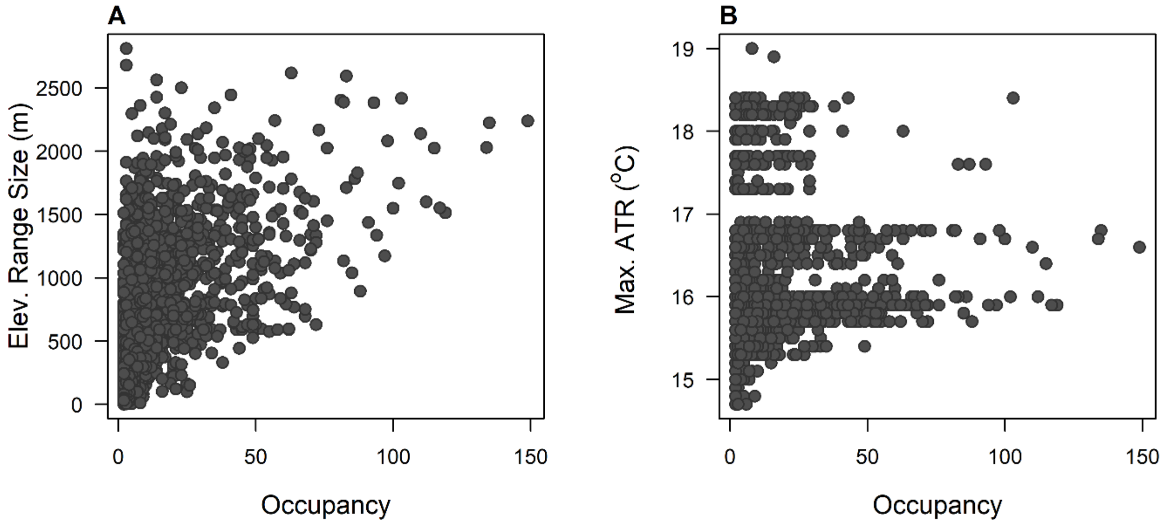

**Figure S8. Multivariate relationships of breadth of elevational distribution with climate variability and other predictors.** These panels show component residual plots whereby the effect of each predictor on elevational range size (left column) and standardized effect sizes (right column) are presented in the context of the other predictors in a regression model. The values in the y-axis correspond to the sum of model residuals plus the "component". The component was calculated as the values predicted by the coefficients of each variable in the model. The component is also shown as the solid black line in each panel. The dashed line shows the trend based on a locally-weighted polynomial regression. These results correspond to the same regressions depicted in Figure 4. The effects of interactions are shown in Figure 5.

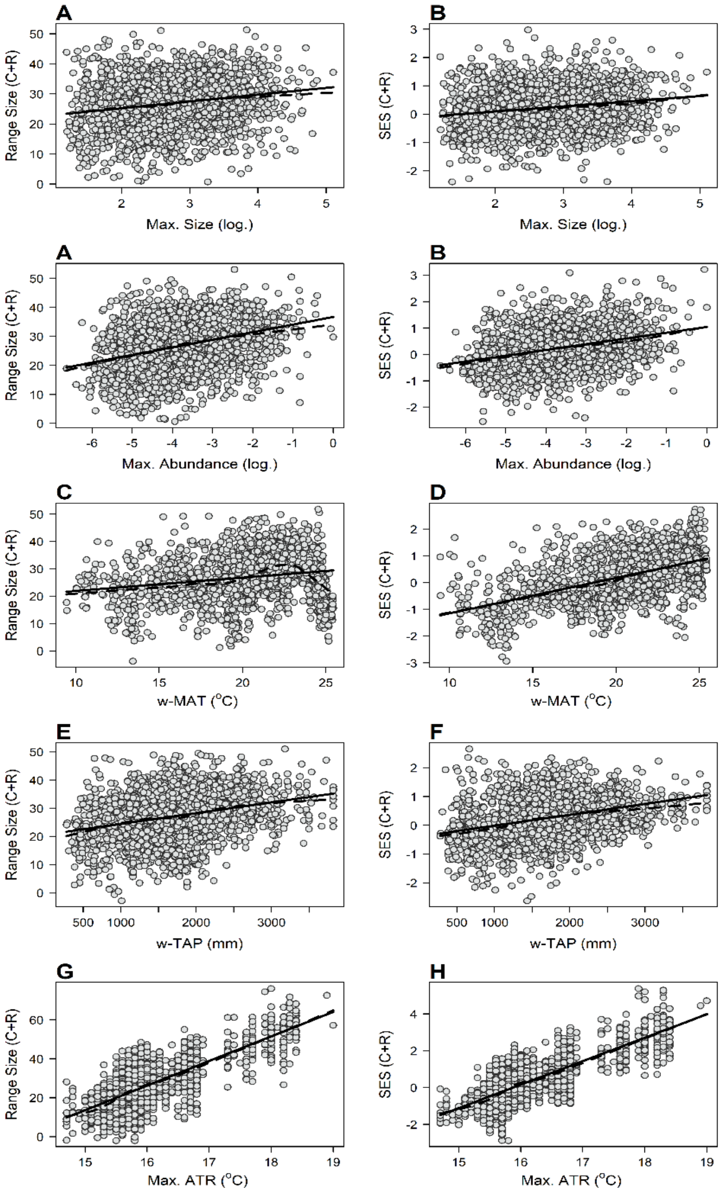

**Figure S9. Effects of climate variability and other predictors on the breadth of elevational distributions – analyses using species present in at least 3 plots or having at least 5 individuals. (A)** Standardized coefficients showing the effect of each predictor on elevational range size. The height of each bar indicates the coefficient estimate, while the lines show the 95% confidence interval. Ives' $R^2_{resid}$ is also shown. **(B)** Same as (A), but where the response variable was the standardized effect sizes of range size. **(C)** Empirical values of range size plotted against predictions made by the regression model in (A). The dashed line shows the trend based on a locally weighted polynomial regression, while the dashed line is the 1:1 correspondence line. **(D)** Same as (C), but where the response variable was the standardized effect sizes of range size (regression model in B). Size: maximum size (90th percentile of diameter at breast height); Abund.: maximum local abundance; w-MAT: abundance-weighted mean annual temperature; w-TAP: abundance-weighted total annual precipitation; ATR: maximum annual temperature range.

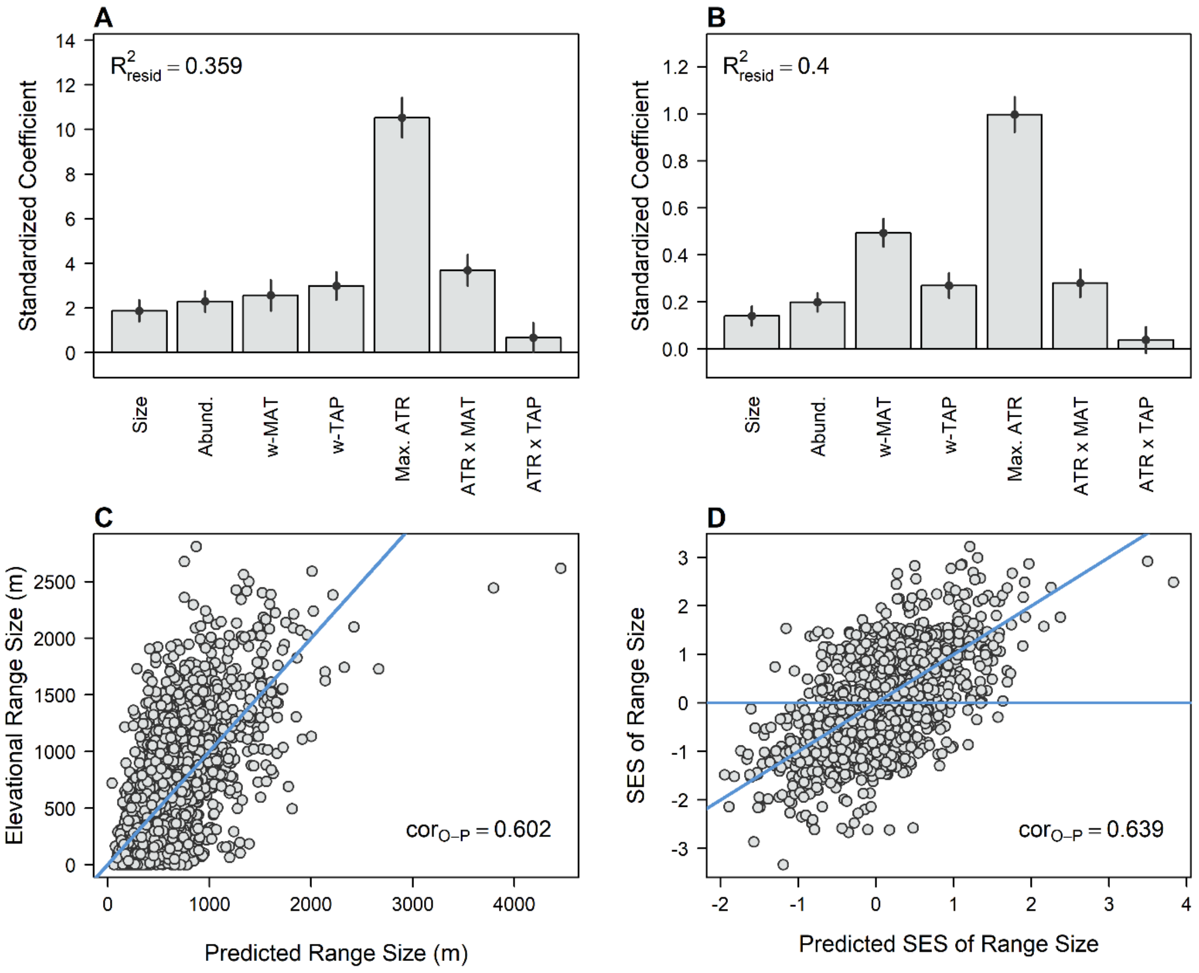

**Figure S10. Effects of climate variability and other predictors on the breadth of elevational distributions – analyses using species present in at least 2 plots among a subset of forest plots that have been selected to standardize sampling effort across elevations. (A)** Standardized coefficients showing the effect of each predictor on elevational range size. The height of each bar indicates the coefficient estimate, while the lines show the 95% confidence interval. Ives' $R^2_{resid}$ is also shown. **(B)** Same as (A), but where the response variable was the standardized effect sizes of range size. **(C)** Empirical values of range size plotted against predictions made by the regression model in (A). The dashed line shows the trend based on a locally-weighted polynomial regression, while the dashed line is the 1:1 correspondence line. **(D)** Same as (C), but where the response variable was the standardized effect sizes of range size (regression model in B). Size: maximum size (90$^{th}$ percentile of diameter at breast height); Abund.: maximum local abundance; w-MAT: abundance-weighted mean annual temperature; w-TAP: abundance-weighted total annual precipitation; ATR: maximum annual temperature range.

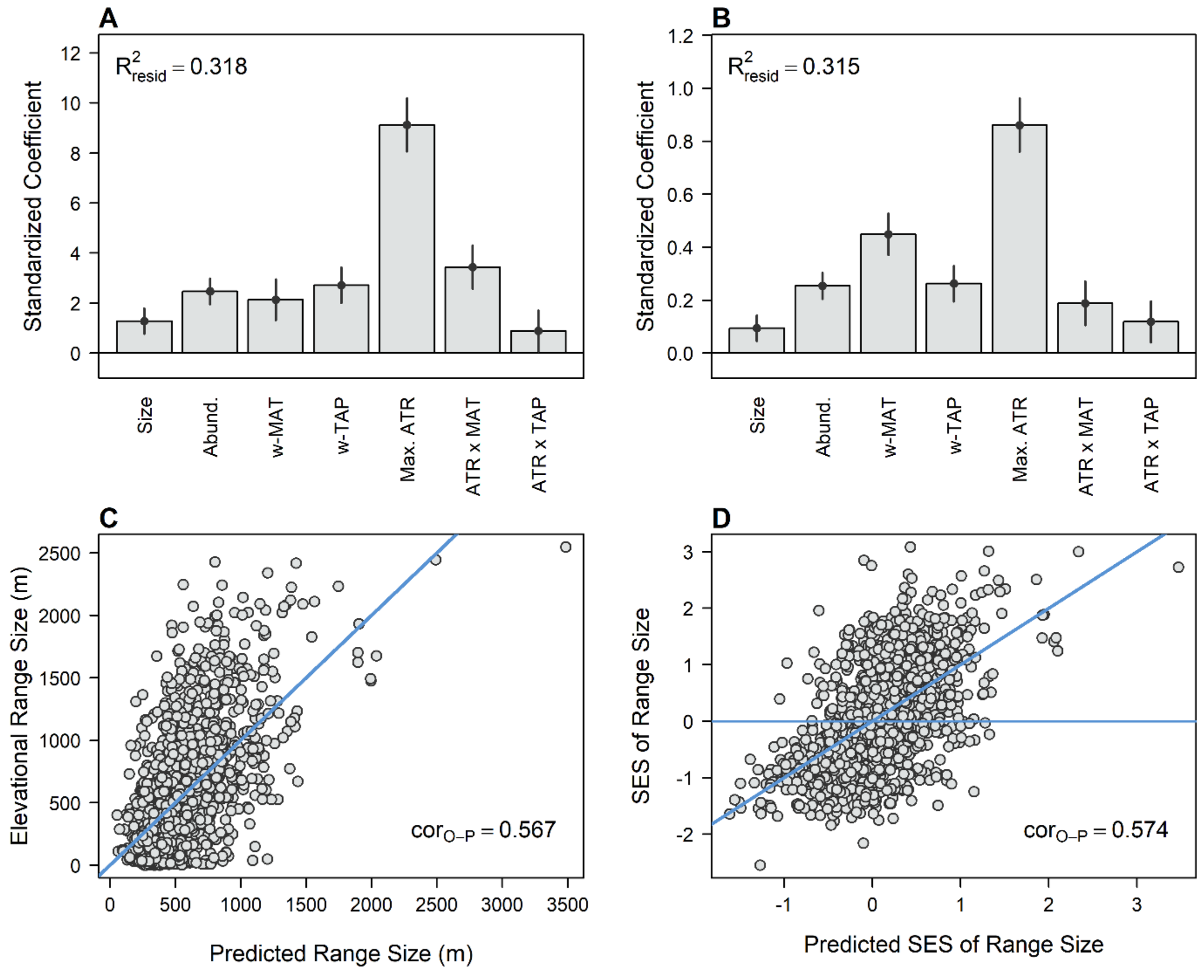

### ADDITIONAL TABLES

**Table S1. Univariate and multivariate models to test for the effects of temperature variability and other predictors on size of elevational distributions.** Phylogenetic generalized least-square regressions (PGLS) were used. Elevational range size (ERS) or a standardized effect size for range size (SES) were modeled as the response variable in separate models. For each response, we created three types of models: univariate models where each predictor was evaluated independently, models only with covariables that did not include the effect of climate variability, and finally models with all covariables, the effect of climate variability and its interactions. Additionally, we used two variables to represent climate variability: maximum diurnal temperature range (max. dtr) and maximum annual temperature range (max. ATR). Covariable predictors included species maximum size, maximum abundance among occupied plots, and abundance-weighted total annual precipitation (w-TAP). For each predictor, we report standardized coefficients, standard deviations, t-values, p-values and variation inflation factors (VIF). PGLS model fit is characterized by Ives' residual $R^2$ value for phylogenetic modes, as well as Pearson's correlation between observed and model-predicted values of the response variable. Finally, we used a likelihood ratio test (LRT) to obtain a model-wide p-value by contrasting each PGLS against a null model. The null model had only an intercept and the same phylogenetic structure estimated for the main PGLS model. For contrast, we report results of corresponding ordinary least-square regressions (OLS), which did not account for phylogenetic relationships among species. For each PGLS and OLS model, we report values of the Akaike information criterion (AIC).

**Table S2. Multivariate models that include the effect of occupancy to evaluate its effects on the importance of other predictors.** Phylogenetic generalized least-square regressions (PGLS) were used. Elevational range size (ERS) or a standardized effect size for range size (SES) were modeled as the response variable in separate models. For each response, we recreated models with all covariables, the effect of climate variability and its interactions. We used maximum annual temperature range (max. ATR) to represent climate variability. Covariable predictors included species maximum size, maximum abundance among occupied plots, and abundance-weighted total annual precipitation (w-TAP). Additionally, we included the logarithm of occupancy (number of plots where the species is present). For each predictor, we report standardized coefficients, standard deviations, t-values, p-values and variation inflation factors (VIF). PGLS model fit is characterized by Ives' residual $R^2$ value for phylogenetic modes, as well as Pearson's correlation between observed and model-predicted values of the response variable. Finally, we used a likelihood ratio test (LRT) to obtain a model-wide p-value by contrasting each PGLS against a null model. The null model had only an intercept and the same phylogenetic structure estimated for the main PGLS model. For contrast, we report results of corresponding ordinary least-square regressions (OLS), which did not account for phylogenetic relationships among species. For each PGLS and OLS model, we report values of the Akaike information criterion (AIC).

**Table S3. Models to test for the effects of temperature variability on size of elevational distributions using species present in at least 3 plots or having at least 5 individuals.** These results represent the same univariate and multivariate analyses reported in Table S1 using one of the alternative datasets. These alternative analyses are used to ensure that results do not depend on the specific subset of species used. Phylogenetic generalized least-square regressions (PGLS) were used. Elevational range size (ERS) or a standardized effect size for range size (SES) were modeled as the response variable in separate models. For each response, we created three types of models: univariate models where each predictor was evaluated independently, models only with covariables that did not include the effect of climate variability, and finally models with all covariables, the effect of climate variability and its interactions. Additionally, we used two variables to represent climate variability: maximum diurnal temperature range (max. dtr) and maximum annual temperature range (max. ATR). Covariable predictors included species maximum size, maximum abundance among occupied plots, and abundance-weighted total annual precipitation (w-TAP). For each predictor, we report standardized coefficients, standard deviations, t-values, p-values and variation inflation factors (VIF). PGLS model fit is characterized by Ives' residual $R^2$ value for phylogenetic modes, as well as Pearson's correlation between observed and model-predicted values of the response variable. Finally, we used a likelihood ratio test (LRT) to obtain a model-wide p-value by contrasting each PGLS against a null model. The null model had only an intercept and the same phylogenetic structure estimated for the main PGLS model. For contrast, we report results of corresponding ordinary least-square regressions (OLS), which did not account for phylogenetic relationships among species. For each PGLS and OLS model, we report values of the Akaike information criterion (AIC).

**Table S4. Models to test for the effects of temperature variability on size of elevational distributions using a subset of the data where sampling along the elevational gradient has been standardized.** These results represent the same univariate and multivariate analyses reported in Table S1 using one of the alternative datasets. The method used to standardize sampling along the elevational gradient is explained in Methods. These alternative analyses are used to ensure that results do not depend on the specific subset of species used. Phylogenetic generalized least-square regressions (PGLS) were used. Elevational range size (ERS) or a standardized effect size for range size (SES) were modeled as the response variable in separate models. For each response, we created three types of models: univariate models where each predictor was evaluated independently, models only with covariables that did not include the effect of climate variability, and finally models with all covariables, the effect of climate variability and its interactions. Additionally, we used two variables to represent climate variability: maximum diurnal temperature range (max. dtr) and maximum annual temperature range (max. ATR). Covariable predictors included species maximum size, maximum abundance among occupied plots, and abundance-weighted total annual precipitation (w-TAP). For each predictor, we report standardized coefficients, standard deviations, t-values, p-values and variation inflation factors (VIF). PGLS model fit is characterized by Ives' residual $R^2$ value for phylogenetic modes, as well as Pearson's correlation between observed and model-predicted values of the response variable. Finally, we used a likelihood ratio test (LRT) to obtain a model-wide p-value by contrasting each PGLS against a null model. The null model had only an intercept and the same phylogenetic structure estimated for the main PGLS model. For contrast, we report results of corresponding ordinary least-square regressions (OLS), which did not account for phylogenetic relationships among species. For each PGLS and OLS model, we report values of the Akaike information criterion (AIC).

| Response | Model | Predictor | Coefficient: PGLS | Std. Error: PGLS | t-Value: PGLS | p-Value: PGLS | VIF: PGLS | AIC: PGLS | cor_O-P: PGLS | R2_resid: PGLS | Likelihood Ratio: PGLS | P: GLS | Coefficient: OLS | Std. Error: OLS | t-Value: OLS | p-Value: OLS | AIC: OLS | Adj. R2: OLS | P: OLS |
|---|---|---|---|---|---|---|---|---|---|---|---|---|---|---|---|---|---|---|---|
| ERS | **Elev.** | Intercept | 24.678 | 0.602 | 41.026 | 0.000 | | 13933.27 | 0.056 | 0.005 | 7.05 | 0.008 | 24.162 | 0.254 | 95.084 | 0.000 | 13938.88 | 0.003 | 0.017 |
| | | Elev. Midpoint | 0.738 | 0.277 | 2.665 | 0.008 | | | | 0.005 | | | 0.609 | 0.254 | 2.394 | 0.017 | | | |
| | **Max. Size** | Intercept | 24.816 | 0.765 | 32.425 | 0.000 | | 13816.12 | 0.250 | 0.072 | 124.20 | 0.000 | 24.162 | 0.246 | 98.038 | 0.000 | 13826.85 | 0.062 | 0.000 |
| | | log(Max. Size) | 3.079 | 0.269 | 11.447 | 0.000 | | | | | | | 2.717 | 0.247 | 11.023 | 0.000 | | | |
| | **Max. Abund.** | Intercept | 24.484 | 0.680 | 36.005 | 0.000 | | 13650.67 | 0.375 | 0.151 | 289.65 | 0.000 | 24.162 | 0.236 | 102.424 | 0.000 | 13666.55 | 0.140 | 0.000 |
| | | log(Max. Abund.) | 4.243 | 0.239 | 17.752 | 0.000 | | | | | | | 4.087 | 0.236 | 17.319 | 0.000 | | | |
| | **w-MAT** | Intercept | 24.709 | 0.625 | 39.536 | 0.000 | | 13926.36 | 0.079 | 0.009 | 13.96 | 0.000 | 24.162 | 0.254 | 95.231 | 0.000 | 13933.21 | 0.006 | 0.001 |
| | | w-MAT | -1.046 | 0.278 | -3.762 | 0.000 | | | | | | | -0.858 | 0.254 | -3.381 | 0.001 | | | |
| | **w-TAP** | Intercept | 24.634 | 0.578 | 42.653 | 0.000 | | 13940.24 | 0.002 | 0.000 | 0.08 | 0.773 | 24.162 | 0.255 | 94.935 | 0.000 | 13944.60 | -0.001 | 0.922 |
| | | w-TAP | -0.074 | 0.258 | -0.289 | 0.773 | | | | | | | -0.025 | 0.255 | -0.098 | 0.922 | | | |
| | **Max. dtr** | Intercept | 25.071 | 1.012 | 24.767 | 0.000 | | 13561.02 | 0.398 | 0.198 | 379.30 | 0.000 | 24.162 | 0.233 | 103.478 | 0.000 | 13629.08 | 0.158 | 0.000 |
| | | Max. dtr | 5.321 | 0.254 | 20.932 | 0.000 | | | | | | | 4.332 | 0.234 | 18.547 | 0.000 | | | |
| | **Max. ATR** | Intercept | 24.815 | 0.804 | 30.883 | 0.000 | | 13762.04 | 0.289 | 0.100 | 178.28 | 0.000 | 24.162 | 0.244 | 99.174 | 0.000 | 13784.65 | 0.083 | 0.000 |
| | | Max. ATR | 3.553 | 0.257 | 13.827 | 0.000 | | | | | | | 3.149 | 0.244 | 12.922 | 0.000 | | | |
| | **Covariables** | Intercept | 24.535 | 0.781 | 31.410 | 0.000 | | 13564.46 | 0.426 | 0.195 | 381.86 | 0.000 | 24.162 | 0.230 | 104.868 | 0.000 | 13583.22 | 0.180 | 0.000 |
| | | log(Max. Abund.) | 3.988 | 0.247 | 16.159 | 0.000 | 1.111 | | | | | | 3.832 | 0.243 | 15.785 | 0.000 | | | |
| | | w-MAT | -0.434 | 0.284 | -1.529 | 0.126 | 1.164 | | | | | | -0.390 | 0.250 | -1.564 | 0.118 | | | |
| | | log(Max. Size) | 2.297 | 0.258 | 8.919 | 0.000 | 1.039 | | | | | | 2.053 | 0.235 | 8.752 | 0.000 | | | |
| | | w-TAP | 0.947 | 0.253 | 3.739 | 0.000 | 1.166 | | | | | | 0.891 | 0.249 | 3.579 | 0.000 | | | |
| | **Covariables x dtr** | Intercept | 26.409 | 0.472 | 55.913 | 0.000 | | 12635.00 | 0.716 | 0.514 | 1317.32 | 0.000 | 26.088 | 0.233 | 111.849 | 0.000 | 12641.83 | 0.510 | 0.000 |
| | | log(Max. | 2.254 | 0.197 | 11.452 | 0.000 | 1.192 | | | | | | 2.174 | 0.194 | 11.203 | 0.000 | | | |
| | | Max. dtr | 11.690 | 0.362 | 32.255 | 0.000 | 3.695 | | | | | | 11.779 | 0.362 | 32.528 | 0.000 | | | |
| | | w-MAT | 8.031 | 0.375 | 21.420 | 0.000 | 3.679 | | | | | | 8.060 | 0.364 | 22.144 | 0.000 | | | |
| | | mat.weighted. | 2.007 | 0.239 | 8.394 | 0.000 | 2.704 | | | | | | 1.981 | 0.239 | 8.303 | 0.000 | | | |
| | | log(Max. Size) | 1.182 | 0.197 | 5.995 | 0.000 | 1.066 | | | | | | 1.100 | 0.184 | 5.989 | 0.000 | | | |
| | | w-TAP | 0.061 | 0.214 | 0.287 | 0.774 | 1.409 | | | | | | 0.031 | 0.211 | 0.148 | 0.882 | | | |
| | | tap.weighted. | 0.748 | 0.313 | 2.391 | 0.017 | 2.909 | | | | | | 0.731 | 0.312 | 2.342 | 0.019 | | | |
| | **Covariables x ATR** | Intercept | 27.011 | 0.616 | 43.838 | 0.000 | | 13061.77 | 0.618 | 0.389 | 890.55 | 0.000 | 26.899 | 0.249 | 108.098 | 0.000 | 13072.88 | 0.380 | 0.000 |
| | | log(Max. | 3.033 | 0.219 | 13.869 | 0.000 | 1.156 | | | | | | 2.928 | 0.215 | 13.626 | 0.000 | | | |
| | | Max. ATR | 9.592 | 0.425 | 22.570 | 0.000 | 4.071 | | | | | | 9.713 | 0.425 | 22.828 | 0.000 | | | |
| | | w-MAT | 1.730 | 0.324 | 5.337 | 0.000 | 2.069 | | | | | | 1.938 | 0.307 | 6.319 | 0.000 | | | |
| | | mat.weighted. | 3.735 | 0.323 | 11.572 | 0.000 | 5.877 | | | | | | 3.776 | 0.324 | 11.648 | 0.000 | | | |
| | | log(Max. Size) | 1.688 | 0.224 | 7.548 | 0.000 | 1.054 | | | | | | 1.536 | 0.205 | 7.482 | 0.000 | | | |
| | | w-TAP | 2.602 | 0.285 | 9.137 | 0.000 | 1.952 | | | | | | 2.500 | 0.281 | 8.906 | 0.000 | | | |
| | | tap.weighted. | 0.309 | 0.304 | 1.016 | 0.310 | 3.591 | | | | | | 0.260 | 0.305 | 0.853 | 0.394 | | | |
| | **Elev.** | Intercept | 0.077 | 0.065 | 1.181 | 0.238 | | 5100.00 | 0.151 | 0.013 | 30.68 | 0.000 | -0.005 | 0.023 | -0.207 | 0.836 | 5108.95 | 0.022 | 0.000 |
| | | Elev. Midpoint | -0.144 | 0.026 | -5.644 | 0.000 | | | | | | | -0.149 | 0.023 | -6.535 | 0.000 | | | |
| | **Max. Size** | Intercept | 0.109 | 0.087 | 1.246 | 0.213 | | 5017.57 | 0.234 | 0.063 | 113.11 | 0.000 | -0.005 | 0.022 | -0.211 | 0.833 | 5047.81 | 0.054 | 0.000 |
| | | log(Max. Size) | 0.269 | 0.025 | 10.855 | 0.000 | | | | | | | 0.231 | 0.022 | 10.309 | 0.000 | | | |
| | **Max. Abund.** | Intercept | 0.075 | 0.086 | 0.871 | 0.384 | | 4897.57 | 0.327 | 0.123 | 233.11 | 0.000 | -0.005 | 0.022 | -0.217 | 0.828 | 4944.78 | 0.106 | 0.000 |
| | | log(Max. | 0.349 | 0.022 | 15.814 | 0.000 | | | | | | | 0.322 | 0.022 | 14.774 | 0.000 | | | |
| | **w-MAT** | Intercept | 0.081 | 0.067 | 1.206 | 0.228 | | 5114.76 | 0.120 | 0.006 | 15.92 | 0.000 | -0.005 | 0.023 | -0.206 | 0.836 | 5124.75 | 0.014 | 0.000 |
| | | w-MAT | 0.105 | 0.026 | 4.080 | 0.000 | | | | | | | 0.118 | 0.023 | 5.160 | 0.000 | | | |
| | **w-TAP** | Intercept | 0.093 | 0.076 | 1.228 | 0.220 | | 5129.38 | 0.037 | 0.000 | 1.30 | 0.254 | -0.005 | 0.023 | -0.205 | 0.837 | 5148.71 | 0.001 | 0.114 |

| | | | | | | | | | | | | | | | | | | | |
|---|---|---|---|---|---|---|---|---|---|---|---|---|---|---|---|---|---|---|---|
| | **w-TAP** | w-TAP | 0.027 | 0.023 | 1.143 | 0.253 | | 5129.38 | 0.037 | 0.000 | 1.30 | 0.254 | 0.036 | 0.023 | 1.581 | 0.114 | 5148.71 | 0.001 | 0.114 |
| | **Max. dtr** | Intercept | 0.120 | 0.104 | 1.159 | 0.247 | | 5008.93 | 0.194 | 0.069 | 121.75 | 0.000 | -0.005 | 0.023 | -0.209 | 0.835 | 5080.89 | 0.037 | 0.000 |
| | | Max. dtr | 0.284 | 0.025 | 11.499 | 0.000 | | | | | | | 0.191 | 0.023 | 8.462 | 0.000 | | | |
| | **Max. ATR** | Intercept | 0.110 | 0.101 | 1.091 | 0.275 | | 4978.10 | 0.235 | 0.084 | 152.58 | 0.000 | -0.005 | 0.022 | -0.211 | 0.833 | 5046.84 | 0.055 | 0.000 |
| | | Max. ATR | 0.305 | 0.024 | 12.830 | 0.000 | | | | | | | 0.232 | 0.022 | 10.358 | 0.000 | | | |
| | **Covariables** | Intercept | 0.071 | 0.086 | 0.820 | 0.412 | | 4765.27 | 0.417 | 0.187 | 371.41 | 0.000 | -0.005 | 0.021 | -0.225 | 0.822 | 4805.40 | 0.173 | 0.000 |
| | | log(Max. | 0.357 | 0.022 | 15.927 | 0.000 | 1.111 | | | | | | 0.339 | 0.022 | 15.370 | 0.000 | | | |
| | | w-MAT | 0.179 | 0.026 | 6.789 | 0.000 | 1.161 | | | | | | 0.172 | 0.023 | 7.567 | 0.000 | | | |
| **SES** | | log(Max. Size) | 0.208 | 0.024 | 8.799 | 0.000 | 1.040 | | | | | | 0.168 | 0.021 | 7.868 | 0.000 | | | |
| | | w-TAP | 0.053 | 0.023 | 2.303 | 0.021 | 1.164 | | | | | | 0.045 | 0.023 | 1.990 | 0.047 | | | |
| | **Covariables x dtr** | Intercept | 0.047 | 0.059 | 0.796 | 0.426 | | 4231.14 | 0.627 | 0.391 | 911.54 | 0.000 | -0.015 | 0.024 | -0.617 | 0.538 | 4248.12 | 0.391 | 0.000 |
| | | log(Max. | 0.223 | 0.020 | 11.202 | 0.000 | 1.192 | | | | | | 0.211 | 0.020 | 10.759 | 0.000 | | | |
| | | Max. dtr | 0.917 | 0.037 | 25.085 | 0.000 | 3.559 | | | | | | 0.937 | 0.037 | 25.605 | 0.000 | | | |
| | | w-MAT | 0.879 | 0.038 | 22.993 | 0.000 | 3.526 | | | | | | 0.891 | 0.037 | 24.233 | 0.000 | | | |
| | | mat.weighted. | -0.060 | 0.024 | -2.479 | 0.013 | 2.657 | | | | | | -0.063 | 0.024 | -2.593 | 0.010 | | | |
| | | log(Max. Size) | 0.117 | 0.020 | 5.754 | 0.000 | 1.068 | | | | | | 0.094 | 0.019 | 5.081 | 0.000 | | | |
| | | w-TAP | 0.073 | 0.022 | 3.366 | 0.001 | 1.404 | | | | | | 0.068 | 0.021 | 3.204 | 0.001 | | | |
| | | tap.weighted. | 0.136 | 0.031 | 4.333 | 0.000 | 2.836 | | | | | | 0.132 | 0.032 | 4.194 | 0.000 | | | |
| | **Covariables x ATR** | Intercept | 0.237 | 0.060 | 3.928 | 0.000 | | 4148.17 | 0.647 | 0.419 | 994.50 | 0.000 | 0.198 | 0.022 | 9.067 | 0.000 | 4166.68 | 0.417 | 0.000 |
| | | log(Max. | 0.252 | 0.019 | 13.129 | 0.000 | 1.157 | | | | | | 0.242 | 0.019 | 12.792 | 0.000 | | | |
| | | Max. ATR | 0.968 | 0.037 | 25.973 | 0.000 | 4.014 | | | | | | 0.989 | 0.037 | 26.448 | 0.000 | | | |
| | | w-MAT | 0.474 | 0.029 | 16.559 | 0.000 | 2.035 | | | | | | 0.493 | 0.027 | 18.299 | 0.000 | | | |
| | | mat.weighted. | 0.280 | 0.028 | 9.891 | 0.000 | 5.806 | | | | | | 0.285 | 0.028 | 10.022 | 0.000 | | | |
| | | log(Max. Size) | 0.136 | 0.020 | 6.877 | 0.000 | 1.055 | | | | | | 0.111 | 0.018 | 6.162 | 0.000 | | | |
| | | w-TAP | 0.257 | 0.025 | 10.300 | 0.000 | 1.946 | | | | | | 0.249 | 0.025 | 10.089 | 0.000 | | | |
| | | tap.weighted. | 0.018 | 0.027 | 0.685 | 0.494 | 3.550 | | | | | | 0.012 | 0.027 | 0.465 | 0.642 | | | |

| Response | Model | Predictor | Coefficient: PGLS | Std. Error: PGLS | t-Value: PGLS | p-Value: PGLS | VIF: PGLS | AIC: PGLS | cor_O-P: PGLS | R2_resid: PGLS | Likelihood Ratio: PGLS | P: GLS | Coefficient: OLS | Std. Error: OLS | t-Value: OLS | p-Value: OLS | AIC: OLS | Adj. R2: OLS | F | P: OLS |
|---|---|---|---|---|---|---|---|---|---|---|---|---|---|---|---|---|---|---|---|---|
| | | Intercept | 25.942 | 0.577 | 44.971 | 0.000 | NA | | | | | | 26.014 | 0.220 | 118.495 | 0.000 | | | | |
| | | log(Max. Abund.) | -0.827 | 0.248 | -3.329 | 0.001 | 1.966 | | | | | | -0.891 | 0.245 | -3.644 | 0.000 | | | | |
| | | Max. ATR | 6.361 | 0.393 | 16.202 | 0.000 | 4.554 | | | | | | 6.473 | 0.394 | 16.444 | 0.000 | | | | |
| | **Covariables** | w-MAT | 0.078 | 0.291 | 0.266 | 0.790 | 2.163 | | | | | | 0.223 | 0.276 | 0.806 | 0.420 | | | | |
| **ERS** | **x ATR +** | mat.weighted.mean:ATR.max | 2.933 | 0.282 | 10.385 | 0.000 | 5.911 | 12550.757 | 0.730 | 0.539 | 1403.561 | 0.000 | 2.988 | 0.284 | 10.526 | 0.000 | 12563.798 | 0.531 | 260.108 | 0.000 |
| | **Occupancy** | occup.n | 6.532 | 0.269 | 24.278 | 0.000 | 2.330 | | | | | | 6.521 | 0.269 | 24.221 | 0.000 | | | | |
| | | log(Max. Size) | 0.247 | 0.205 | 1.209 | 0.227 | 1.154 | | | | | | 0.038 | 0.189 | 0.202 | 0.840 | | | | |
| | | w-TAP | 1.047 | 0.256 | 4.089 | 0.000 | 2.079 | | | | | | 0.998 | 0.252 | 3.960 | 0.000 | | | | |
| | | tap.weighted.mean:ATR.max | -0.345 | 0.265 | -1.299 | 0.194 | 3.601 | | | | | | -0.368 | 0.267 | -1.381 | 0.167 | | | | |
| | | Intercept | 0.153 | 0.053 | 2.895 | 0.004 | NA | | | | | | 0.131 | 0.020 | 6.527 | 0.000 | | | | |
| | | log(Max. Abund.) | -0.041 | 0.023 | -1.826 | 0.068 | 1.966 | | | | | | -0.050 | 0.022 | -2.227 | 0.026 | | | | |
| | | Max. ATR | 0.724 | 0.036 | 20.229 | 0.000 | 4.549 | | | | | | 0.742 | 0.036 | 20.641 | 0.000 | | | | |
| | **Covariables** | w-MAT | 0.350 | 0.027 | 13.194 | 0.000 | 2.160 | | | | | | 0.362 | 0.025 | 14.377 | 0.000 | | | | |
| **SES** | **x ATR +** | mat.weighted.mean:ATR.max | 0.219 | 0.026 | 8.506 | 0.000 | 5.906 | 3780.548 | 0.725 | 0.525 | 1364.131 | 0.000 | 0.225 | 0.026 | 8.697 | 0.000 | 3797.616 | 0.524 | 252.911 | 0.000 |
| | **Occupancy** | occup.n | 0.495 | 0.025 | 20.189 | 0.000 | 2.329 | | | | | | 0.497 | 0.025 | 20.232 | 0.000 | | | | |
| | | log(Max. Size) | 0.024 | 0.019 | 1.301 | 0.193 | 1.154 | | | | | | -0.003 | 0.017 | -0.178 | 0.859 | | | | |
| | | w-TAP | 0.139 | 0.023 | 5.962 | 0.000 | 2.079 | | | | | | 0.134 | 0.023 | 5.838 | 0.000 | | | | |
| | | tap.weighted.mean:ATR.max | -0.031 | 0.024 | -1.290 | 0.197 | 3.598 | | | | | | -0.035 | 0.024 | -1.457 | 0.145 | | | | |

| Response | Model | Predictor | Coefficient: PGLS | Std. Error: PGLS | t-Value: PGLS | p-Value: PGLS | VIF: PGLS | AIC: PGLS | cor_O-P: PGLS | R2_resid : PGLS | Likelihood Ratio: PGLS | P: GLS | Coefficient: OLS | Std. Error: OLS | t-Value: OLS | p-Value: OLS | AIC: OLS | Adj. R2: OLS | F | P: OLS |
|---|---|---|---|---|---|---|---|---|---|---|---|---|---|---|---|---|---|---|---|---|
| ERS | Elev. | Intercept | 25.278 | 0.729 | 34.673 | 0 | NA | 13158.60 | -0.002 | 0 | 0.13 | 0.719 | 24.463 | 0.273 | 89.629 | 0 | 13171.58 | -0.001 | 0.011 | 0.918 |
| ERS | Elev. | Elev. Midpoint | 0.109 | 0.302 | 0.361 | 0.718 | NA | 13158.60 | -0.002 | 0 | 0.13 | 0.719 | -0.028 | 0.273 | -0.103 | 0.918 | 13171.58 | -0.001 | 0.011 | 0.918 |
| ERS | Max. Size | Intercept | 25.316 | 0.782 | 32.362 | 0 | NA | 13059.29 | 0.241 | 0.059 | 99.443 | 0 | 24.463 | 0.265 | 92.343 | 0 | 13069.39 | 0.057 | 105.188 | 0 |
| ERS | Max. Size | log(Max. Size) | 2.932 | 0.289 | 10.132 | 0 | NA | 13059.29 | 0.241 | 0.059 | 99.443 | 0 | 2.718 | 0.265 | 10.256 | 0 | 13069.39 | 0.057 | 105.188 | 0 |
| ERS | Max. Abund. | Intercept | 25.061 | 0.743 | 33.743 | 0 | NA | 13019.40 | 0.274 | 0.079 | 139.332 | 0 | 24.463 | 0.262 | 93.2 | 0 | 13037.71 | 0.075 | 139.084 | 0 |
| ERS | Max. Abund. | log(Max. Abund.) | 3.206 | 0.266 | 12.047 | 0 | NA | 13019.40 | 0.274 | 0.079 | 139.332 | 0 | 3.096 | 0.263 | 11.793 | 0 | 13037.71 | 0.075 | 139.084 | 0 |
| ERS | w-MAT | Intercept | 25.305 | 0.744 | 34.005 | 0 | NA | 13156.46 | 0.022 | 0.002 | 2.269 | 0.132 | 24.463 | 0.273 | 89.651 | 0 | 13170.73 | 0 | 0.859 | 0.354 |
| ERS | w-MAT | w-MAT | -0.457 | 0.302 | -1.513 | 0.13 | NA | 13156.46 | 0.022 | 0.002 | 2.269 | 0.132 | -0.253 | 0.273 | -0.927 | 0.354 | 13170.73 | 0 | 0.859 | 0.354 |
| ERS | w-TAP | Intercept | 25.267 | 0.725 | 34.874 | 0 | NA | 13158.41 | 0.017 | 0 | 0.324 | 0.569 | 24.463 | 0.273 | 89.642 | 0 | 13171.08 | 0 | 0.51 | 0.475 |
| ERS | w-TAP | w-TAP | 0.157 | 0.277 | 0.569 | 0.569 | NA | 13158.41 | 0.017 | 0 | 0.324 | 0.569 | 0.195 | 0.273 | 0.714 | 0.475 | 13171.08 | 0 | 0.51 | 0.475 |
| ERS | Max. dtr | Intercept | 25.616 | 1.099 | 23.302 | 0 | NA | 12871.70 | 0.353 | 0.162 | 287.033 | 0 | 24.463 | 0.255 | 95.802 | 0 | 12943.41 | 0.124 | 243.795 | 0 |
| ERS | Max. dtr | Max. dtr | 4.973 | 0.276 | 18.007 | 0 | NA | 12871.70 | 0.353 | 0.162 | 287.033 | 0 | 3.988 | 0.255 | 15.614 | 0 | 12943.41 | 0.124 | 243.795 | 0 |
| ERS | Max. ATR | Intercept | 25.334 | 0.886 | 28.588 | 0 | NA | 13008.76 | 0.271 | 0.089 | 149.973 | 0 | 24.463 | 0.263 | 93.108 | 0 | 13041.11 | 0.073 | 135.415 | 0 |
| ERS | Max. ATR | Max. ATR | 3.477 | 0.276 | 12.605 | 0 | NA | 13008.76 | 0.271 | 0.089 | 149.973 | 0 | 3.058 | 0.263 | 11.637 | 0 | 13041.11 | 0.073 | 135.415 | 0 |
| ERS | Covariables | Intercept | 25.07 | 0.791 | 31.712 | 0 | NA | 12935.43 | 0.354 | 0.128 | 229.304 | 0 | 24.463 | 0.256 | 95.739 | 0 | 12948.64 | 0.123 | 61.06 | 0 |
| ERS | Covariables | log(Max. Abund.) | 3.13 | 0.273 | 11.465 | 0 | 1.102 | 12935.43 | 0.354 | 0.128 | 229.304 | 0 | 3.044 | 0.269 | 11.328 | 0 | 12948.64 | 0.123 | 61.06 | 0 |
| ERS | Covariables | w-MAT | 0.067 | 0.314 | 0.212 | 0.832 | 1.191 | 12935.43 | 0.354 | 0.128 | 229.304 | 0 | 0.114 | 0.279 | 0.407 | 0.684 | 12948.64 | 0.123 | 61.06 | 0 |
| ERS | Covariables | log(Max. Size) | 2.525 | 0.282 | 8.94 | 0 | 1.018 | 12935.43 | 0.354 | 0.128 | 229.304 | 0 | 2.356 | 0.258 | 9.14 | 0 | 12948.64 | 0.123 | 61.06 | 0 |
| ERS | Covariables | w-TAP | 0.867 | 0.283 | 3.059 | 0.002 | 1.19 | 12935.43 | 0.354 | 0.128 | 229.304 | 0 | 0.799 | 0.278 | 2.869 | 0.004 | 12948.64 | 0.123 | 61.06 | 0 |
| ERS | Covariables x dtr | Intercept | 26.736 | 0.547 | 48.911 | 0 | NA | 12083.76 | 0.686 | 0.47 | 1086.976 | 0 | 26.163 | 0.258 | 101.226 | 0 | 12094.36 | 0.468 | 216.45 | 0 |
| ERS | Covariables x dtr | log(Max. Abund.) | 1.638 | 0.217 | 7.553 | 0 | 1.155 | 12083.76 | 0.686 | 0.47 | 1086.976 | 0 | 1.603 | 0.214 | 7.492 | 0 | 12094.36 | 0.468 | 216.45 | 0 |
| ERS | Covariables x dtr | Max. dtr | 12.192 | 0.387 | 31.484 | 0 | 3.383 | 12083.76 | 0.686 | 0.47 | 1086.976 | 0 | 12.253 | 0.388 | 31.616 | 0 | 12094.36 | 0.468 | 216.45 | 0 |
| ERS | Covariables x dtr | w-MAT | 8.746 | 0.408 | 21.451 | 0 | 3.462 | 12083.76 | 0.686 | 0.47 | 1086.976 | 0 | 8.752 | 0.394 | 22.226 | 0 | 12094.36 | 0.468 | 216.45 | 0 |
| ERS | Covariables x dtr | mat.weighted. mean:dtr.max | 1.537 | 0.266 | 5.777 | 0 | 2.629 | 12083.76 | 0.686 | 0.47 | 1086.976 | 0 | 1.459 | 0.265 | 5.498 | 0 | 12094.36 | 0.468 | 216.45 | 0 |
| ERS | Covariables x dtr | log(Max. Size) | 1.477 | 0.219 | 6.732 | 0 | 1.043 | 12083.76 | 0.686 | 0.47 | 1086.976 | 0 | 1.383 | 0.203 | 6.8 | 0 | 12094.36 | 0.468 | 216.45 | 0 |
| ERS | Covariables x dtr | w-TAP | 0.221 | 0.242 | 0.912 | 0.362 | 1.445 | 12083.76 | 0.686 | 0.47 | 1086.976 | 0 | 0.194 | 0.239 | 0.813 | 0.416 | 12094.36 | 0.468 | 216.45 | 0 |
| ERS | Covariables x dtr | tap.weighted. mean:dtr.max | 1.368 | 0.349 | 3.918 | 0 | 2.851 | 12083.76 | 0.686 | 0.47 | 1086.976 | 0 | 1.397 | 0.349 | 4.006 | 0 | 12094.36 | 0.468 | 216.45 | 0 |
| ERS | Covariables x ATR | Intercept | 27.601 | 0.562 | 49.114 | 0 | NA | 12405.66 | 0.602 | 0.359 | 765.072 | 0 | 27.357 | 0.271 | 100.781 | 0 | 12411.72 | 0.36 | 138.652 | 0 |
| ERS | Covariables x ATR | log(Max. Abund.) | 2.296 | 0.235 | 9.755 | 0 | 1.131 | 12405.66 | 0.602 | 0.359 | 765.072 | 0 | 2.257 | 0.232 | 9.718 | 0 | 12411.72 | 0.36 | 138.652 | 0 |
| ERS | Covariables x ATR | Max. ATR | 10.532 | 0.45 | 23.388 | 0 | 3.947 | 12405.66 | 0.602 | 0.359 | 765.072 | 0 | 10.604 | 0.45 | 23.547 | 0 | 12411.72 | 0.36 | 138.652 | 0 |
| ERS | Covariables x ATR | w-MAT | 2.573 | 0.35 | 7.356 | 0 | 2.143 | 12405.66 | 0.602 | 0.359 | 765.072 | 0 | 2.761 | 0.334 | 8.272 | 0 | 12411.72 | 0.36 | 138.652 | 0 |
| ERS | Covariables x ATR | mat.weighted. mean:ATR.max | 3.694 | 0.355 | 10.415 | 0 | 5.8 | 12405.66 | 0.602 | 0.359 | 765.072 | 0 | 3.706 | 0.355 | 10.428 | 0 | 12411.72 | 0.36 | 138.652 | 0 |
| ERS | Covariables x ATR | log(Max. Size) | 1.875 | 0.238 | 7.864 | 0 | 1.032 | 12405.66 | 0.602 | 0.359 | 765.072 | 0 | 1.797 | 0.222 | 8.103 | 0 | 12411.72 | 0.36 | 138.652 | 0 |
| ERS | Covariables x ATR | w-TAP | 2.992 | 0.313 | 9.575 | 0 | 1.995 | 12405.66 | 0.602 | 0.359 | 765.072 | 0 | 2.889 | 0.308 | 9.365 | 0 | 12411.72 | 0.36 | 138.652 | 0 |
| ERS | Covariables x ATR | tap.weighted. mean:ATR.max | 0.678 | 0.335 | 2.024 | 0.043 | 3.628 | 12405.66 | 0.602 | 0.359 | 765.072 | 0 | 0.654 | 0.335 | 1.951 | 0.051 | 12411.72 | 0.36 | 138.652 | 0 |
| SES | Elev. | Intercept | 0.107 | 0.079 | 1.344 | 0.179 | NA | 4756.93 | 0.174 | 0.021 | 38.866 | 0 | -0.006 | 0.024 | -0.257 | 0.797 | 4778.73 | 0.03 | 53.434 | 0 |
| SES | Elev. | Elev. Midpoint | -0.169 | 0.027 | -6.304 | 0 | NA | 4756.93 | 0.174 | 0.021 | 38.866 | 0 | -0.172 | 0.024 | -7.31 | 0 | 4778.73 | 0.03 | 53.434 | 0 |
| SES | Max. Size | Intercept | 0.13 | 0.095 | 1.373 | 0.17 | NA | 4711.27 | 0.216 | 0.05 | 84.525 | 0 | -0.006 | 0.023 | -0.259 | 0.796 | 4749.23 | 0.046 | 84.092 | 0 |
| SES | Max. Size | log(Max. Size) | 0.243 | 0.026 | 9.332 | 0 | NA | 4711.27 | 0.216 | 0.05 | 84.525 | 0 | 0.214 | 0.023 | 9.17 | 0 | 4749.23 | 0.046 | 84.092 | 0 |

| | | | | | | | | | | | | | | | | | | | | |
|---|---|---|---|---|---|---|---|---|---|---|---|---|---|---|---|---|---|---|---|---|
| SES | Max. Abund. | Intercept | 0.099 | 0.093 | 1.063 | 0.288 | NA | 4678.54 | 0.24 | 0.067 | 117.251 | 0 | -0.006 | 0.023 | -0.261 | 0.794 | 4729.39 | 0.057 | 104.994 | 0 |
| SES | Max. Abund. | log(Max. Abund.) | 0.259 | 0.024 | 11.029 | 0 | NA | 4678.54 | 0.24 | 0.067 | 117.251 | 0 | 0.238 | 0.023 | 10.247 | 0 | 4729.39 | 0.057 | 104.994 | 0 |
| SES | w-MAT | Intercept | 0.111 | 0.08 | 1.376 | 0.169 | NA | 4773.91 | 0.142 | 0.011 | 21.884 | 0 | -0.006 | 0.024 | -0.256 | 0.798 | 4796.54 | 0.02 | 35.191 | 0 |
| SES | w-MAT | w-MAT | 0.128 | 0.027 | 4.733 | 0 | NA | 4773.91 | 0.142 | 0.011 | 21.884 | 0 | 0.141 | 0.024 | 5.932 | 0 | 4796.54 | 0.02 | 35.191 | 0 |
| SES | w-TAP | Intercept | 0.121 | 0.087 | 1.391 | 0.164 | NA | 4792.89 | 0.051 | 0.001 | 2.902 | 0.088 | -0.006 | 0.024 | -0.253 | 0.8 | 4826.88 | 0.002 | 4.536 | 0.033 |
| SES | w-TAP | w-TAP | 0.041 | 0.024 | 1.705 | 0.088 | NA | 4792.89 | 0.051 | 0.001 | 2.902 | 0.088 | 0.051 | 0.024 | 2.13 | 0.033 | 4826.88 | 0.002 | 4.536 | 0.033 |
| SES | Max. dtr | Intercept | 0.14 | 0.11 | 1.266 | 0.206 | NA | 4692.25 | 0.182 | 0.061 | 103.547 | 0 | -0.006 | 0.024 | -0.257 | 0.797 | 4773.50 | 0.033 | 58.832 | 0 |
| SES | Max. dtr | Max. dtr | 0.268 | 0.026 | 10.524 | 0 | NA | 4692.25 | 0.182 | 0.061 | 103.547 | 0 | 0.18 | 0.024 | 7.67 | 0 | 4773.50 | 0.033 | 58.832 | 0 |
| SES | Max. ATR | Intercept | 0.123 | 0.106 | 1.157 | 0.248 | NA | 4664.37 | 0.226 | 0.076 | 131.424 | 0 | -0.006 | 0.023 | -0.26 | 0.795 | 4741.75 | 0.05 | 91.945 | 0 |
| SES | Max. ATR | Max. ATR | 0.29 | 0.025 | 11.815 | 0 | NA | 4664.37 | 0.226 | 0.076 | 131.424 | 0 | 0.224 | 0.023 | 9.589 | 0 | 4741.75 | 0.05 | 91.945 | 0 |
| SES | Covariables | Intercept | 0.091 | 0.092 | 0.983 | 0.326 | NA | 4552.81 | 0.365 | 0.137 | 248.985 | 0 | -0.006 | 0.022 | -0.272 | 0.786 | 4591.73 | 0.132 | 65.849 | 0 |
| SES | Covariables | log(Max. Abund.) | 0.284 | 0.024 | 11.955 | 0 | 1.1 | 4552.81 | 0.365 | 0.137 | 248.985 | 0 | 0.272 | 0.023 | 11.609 | 0 | 4591.73 | 0.132 | 65.849 | 0 |
| SES | Covariables | w-MAT | 0.194 | 0.028 | 6.866 | 0 | 1.187 | 4552.81 | 0.365 | 0.137 | 248.985 | 0 | 0.185 | 0.024 | 7.61 | 0 | 4591.73 | 0.132 | 65.849 | 0 |
| SES | Covariables | log(Max. Size) | 0.214 | 0.025 | 8.526 | 0 | 1.018 | 4552.81 | 0.365 | 0.137 | 248.985 | 0 | 0.177 | 0.022 | 7.89 | 0 | 4591.73 | 0.132 | 65.849 | 0 |
| SES | Covariables | w-TAP | 0.045 | 0.025 | 1.84 | 0.066 | 1.186 | 4552.81 | 0.365 | 0.137 | 248.985 | 0 | 0.041 | 0.024 | 1.704 | 0.089 | 4591.73 | 0.132 | 65.849 | 0 |
| SES | Covariables x dtr | Intercept | 0.07 | 0.064 | 1.089 | 0.276 | NA | 4006.61 | 0.615 | 0.372 | 801.183 | 0 | -0.012 | 0.025 | -0.478 | 0.633 | 4026.78 | 0.377 | 148.771 | 0 |
| SES | Covariables x dtr | log(Max. Abund.) | 0.17 | 0.021 | 8.252 | 0 | 1.154 | 4006.61 | 0.615 | 0.372 | 801.183 | 0 | 0.162 | 0.02 | 7.968 | 0 | 4026.78 | 0.377 | 148.771 | 0 |
| SES | Covariables x dtr | Max. dtr | 0.935 | 0.037 | 25.509 | 0 | 3.261 | 4006.61 | 0.615 | 0.372 | 801.183 | 0 | 0.954 | 0.037 | 25.935 | 0 | 4026.78 | 0.377 | 148.771 | 0 |
| SES | Covariables x dtr | w-MAT | 0.892 | 0.039 | 22.833 | 0 | 3.326 | 4006.61 | 0.615 | 0.372 | 801.183 | 0 | 0.899 | 0.037 | 24.051 | 0 | 4026.78 | 0.377 | 148.771 | 0 |
| SES | Covariables x dtr | mat.weighted. mean:dtr.max | -0.062 | 0.025 | -2.453 | 0.014 | 2.587 | 4006.61 | 0.615 | 0.372 | 801.183 | 0 | -0.069 | 0.025 | -2.75 | 0.006 | 4026.78 | 0.377 | 148.771 | 0 |
| SES | Covariables x dtr | log(Max. Size) | 0.124 | 0.021 | 5.844 | 0 | 1.044 | 4006.61 | 0.615 | 0.372 | 801.183 | 0 | 0.102 | 0.019 | 5.262 | 0 | 4026.78 | 0.377 | 148.771 | 0 |
| SES | Covariables x dtr | w-TAP | 0.072 | 0.023 | 3.14 | 0.002 | 1.441 | 4006.61 | 0.615 | 0.372 | 801.183 | 0 | 0.072 | 0.023 | 3.19 | 0.001 | 4026.78 | 0.377 | 148.771 | 0 |
| SES | Covariables x dtr | tap.weighted. mean:dtr.max | 0.154 | 0.033 | 4.654 | 0 | 2.783 | 4006.61 | 0.615 | 0.372 | 801.183 | 0 | 0.156 | 0.033 | 4.714 | 0 | 4026.78 | 0.377 | 148.771 | 0 |
| SES | Covariables x ATR | Intercept | 0.255 | 0.06 | 4.242 | 0 | NA | 3926.39 | 0.639 | 0.4 | 881.407 | 0 | 0.208 | 0.023 | 9.061 | 0 | 3943.21 | 0.406 | 168.386 | 0 |
| SES | Covariables x ATR | log(Max. Abund.) | 0.199 | 0.02 | 9.968 | 0 | 1.13 | 3926.39 | 0.639 | 0.4 | 881.407 | 0 | 0.192 | 0.02 | 9.78 | 0 | 3943.21 | 0.406 | 168.386 | 0 |
| SES | Covariables x ATR | Max. ATR | 0.996 | 0.038 | 26.277 | 0 | 3.832 | 3926.39 | 0.639 | 0.4 | 881.407 | 0 | 1.015 | 0.038 | 26.697 | 0 | 3943.21 | 0.406 | 168.386 | 0 |
| SES | Covariables x ATR | w-MAT | 0.493 | 0.03 | 16.464 | 0 | 2.069 | 3926.39 | 0.639 | 0.4 | 881.407 | 0 | 0.51 | 0.028 | 18.095 | 0 | 3943.21 | 0.406 | 168.386 | 0 |
| SES | Covariables x ATR | mat.weighted. mean:ATR.max | 0.28 | 0.03 | 9.377 | 0 | 5.659 | 3926.39 | 0.639 | 0.4 | 881.407 | 0 | 0.284 | 0.03 | 9.457 | 0 | 3943.21 | 0.406 | 168.386 | 0 |
| SES | Covariables x ATR | log(Max. Size) | 0.14 | 0.021 | 6.822 | 0 | 1.033 | 3926.39 | 0.639 | 0.4 | 881.407 | 0 | 0.12 | 0.019 | 6.388 | 0 | 3943.21 | 0.406 | 168.386 | 0 |
| SES | Covariables x ATR | w-TAP | 0.269 | 0.026 | 10.203 | 0 | 1.982 | 3926.39 | 0.639 | 0.4 | 881.407 | 0 | 0.264 | 0.026 | 10.118 | 0 | 3943.21 | 0.406 | 168.386 | 0 |
| SES | Covariables x ATR | tap.weighted. mean:ATR.max | 0.038 | 0.028 | 1.348 | 0.178 | 3.543 | 3926.39 | 0.639 | 0.4 | 881.407 | 0 | 0.036 | 0.028 | 1.263 | 0.207 | 3943.21 | 0.406 | 168.386 | 0 |

| Response | Model | Predictor | Coefficient: PGLS | Std. Error: PGLS | t-Value: PGLS | p-Value: PGLS | VIF: PGLS | AIC: PGLS | cor_O-P: PGLS | R2_resid: PGLS | Likelihood Ratio: PGLS | P: GLS | Coefficient: OLS | Std. Error: OLS | t-Value: OLS | p-Value: OLS | AIC: OLS | Adj. R2: OLS | F | P: OLS |
|---|---|---|---|---|---|---|---|---|---|---|---|---|---|---|---|---|---|---|---|---|
| **ERS** | **Elev.** | Intercept | 23.794 | 0.698 | 34.112 | 0 | NA | 9582.68 | 0.058 | 0.007 | 6.752 | 0.009 | 23.072 | 0.286 | 80.696 | 0 | 9589.08 | 0.003 | 4.325 | 0.038 |
| **ERS** | **Elev.** | Elev. Midpoint | 0.829 | 0.317 | 2.617 | 0.009 | NA | 9582.68 | 0.058 | 0.007 | 6.752 | 0.009 | 0.595 | 0.286 | 2.08 | 0.038 | 9589.08 | 0.003 | 4.325 | 0.038 |
| **ERS** | **Max. Size** | Intercept | 23.794 | 0.725 | 32.825 | 0 | NA | 9519.59 | 0.23 | 0.056 | 69.843 | 0 | 23.072 | 0.279 | 82.788 | 0 | 9523.56 | 0.052 | 71.673 | 0 |
| **ERS** | **Max. Size** | log(Max. Size) | 2.531 | 0.298 | 8.489 | 0 | NA | 9519.59 | 0.23 | 0.056 | 69.843 | 0 | 2.36 | 0.279 | 8.466 | 0 | 9523.56 | 0.052 | 71.673 | 0 |
| **ERS** | **Max. Abund.** | Intercept | 23.455 | 0.624 | 37.56 | 0 | NA | 9435.10 | 0.335 | 0.114 | 154.333 | 0 | 23.072 | 0.27 | 85.495 | 0 | 9441.19 | 0.111 | 161.375 | 0 |
| **ERS** | **Max. Abund.** | log(Max. Abund.) | 3.488 | 0.273 | 12.797 | 0 | NA | 9435.10 | 0.335 | 0.114 | 154.333 | 0 | 3.429 | 0.27 | 12.703 | 0 | 9441.19 | 0.111 | 161.375 | 0 |
| **ERS** | **w-MAT** | Intercept | 23.878 | 0.734 | 32.534 | 0 | NA | 9572.21 | 0.097 | 0.016 | 17.218 | 0 | 23.072 | 0.285 | 80.945 | 0 | 9581.22 | 0.009 | 12.223 | 0 |
| **ERS** | **w-MAT** | w-MAT | -1.332 | 0.317 | -4.199 | 0 | NA | 9572.21 | 0.097 | 0.016 | 17.218 | 0 | -0.997 | 0.285 | -3.496 | 0 | 9581.22 | 0.009 | 12.223 | 0 |
| **ERS** | **w-TAP** | Intercept | 23.689 | 0.667 | 35.506 | 0 | NA | 9588.61 | 0.016 | 0.001 | 0.816 | 0.366 | 23.072 | 0.286 | 80.57 | 0 | 9593.07 | -0.001 | 0.326 | 0.568 |
| **ERS** | **w-TAP** | w-TAP | -0.265 | 0.292 | -0.906 | 0.365 | NA | 9588.61 | 0.016 | 0.001 | 0.816 | 0.366 | -0.164 | 0.286 | -0.571 | 0.568 | 9593.07 | -0.001 | 0.326 | 0.568 |
| **ERS** | **Max. dtr** | Intercept | 24.491 | 0.974 | 25.143 | 0 | NA | 9370.71 | 0.356 | 0.166 | 218.716 | 0 | 23.072 | 0.268 | 86.212 | 0 | 9419.82 | 0.126 | 185.608 | 0 |
| **ERS** | **Max. dtr** | Max. dtr | 4.62 | 0.294 | 15.709 | 0 | NA | 9370.71 | 0.356 | 0.166 | 218.716 | 0 | 3.647 | 0.268 | 13.624 | 0 | 9419.82 | 0.126 | 185.608 | 0 |
| **ERS** | **Max. ATR** | Intercept | 23.988 | 0.825 | 29.076 | 0 | NA | 9477.36 | 0.268 | 0.09 | 112.072 | 0 | 23.072 | 0.276 | 83.63 | 0 | 9497.66 | 0.071 | 99.255 | 0 |
| **ERS** | **Max. ATR** | Max. ATR | 3.2 | 0.293 | 10.914 | 0 | NA | 9477.36 | 0.268 | 0.09 | 112.072 | 0 | 2.75 | 0.276 | 9.963 | 0 | 9497.66 | 0.071 | 99.255 | 0 |
| **ERS** | **Covariables** | Intercept | 23.485 | 0.625 | 37.596 | 0 | NA | 9393.15 | 0.384 | 0.147 | 202.277 | 0 | 23.072 | 0.265 | 87.14 | 0 | 9395.41 | 0.145 | 55.069 | 0 |
| **ERS** | **Covariables** | log(Max. Abund.) | 3.22 | 0.289 | 11.128 | 0 | 1.167 | 9393.15 | 0.384 | 0.147 | 202.277 | 0 | 3.205 | 0.286 | 11.222 | 0 | 9395.41 | 0.145 | 55.069 | 0 |
| **ERS** | **Covariables** | w-MAT | -0.568 | 0.33 | -1.722 | 0.085 | 1.282 | 9393.15 | 0.384 | 0.147 | 202.277 | 0 | -0.432 | 0.302 | -1.432 | 0.152 | 9395.41 | 0.145 | 55.069 | 0 |
| **ERS** | **Covariables** | log(Max. Size) | 1.783 | 0.288 | 6.191 | 0 | 1.047 | 9393.15 | 0.384 | 0.147 | 202.277 | 0 | 1.768 | 0.27 | 6.551 | 0 | 9395.41 | 0.145 | 55.069 | 0 |
| **ERS** | **Covariables** | w-TAP | 0.895 | 0.305 | 2.935 | 0.003 | 1.268 | 9393.15 | 0.384 | 0.147 | 202.277 | 0 | 0.866 | 0.299 | 2.898 | 0.004 | 9395.41 | 0.145 | 55.069 | 0 |
| **ERS** | **Covariables x dtr** | Intercept | 25.805 | 0.442 | 58.414 | 0 | NA | 8812.77 | 0.679 | 0.457 | 788.657 | 0 | 25.378 | 0.287 | 88.282 | 0 | 8813.50 | 0.458 | 155.631 | 0 |
| **ERS** | **Covariables x dtr** | log(Max. Abund.) | 2.252 | 0.234 | 9.632 | 0 | 1.213 | 8812.77 | 0.679 | 0.457 | 788.657 | 0 | 2.275 | 0.232 | 9.816 | 0 | 8813.50 | 0.458 | 155.631 | 0 |
| **ERS** | **Covariables x dtr** | Max. dtr | 11.974 | 0.472 | 25.365 | 0 | 4.595 | 8812.77 | 0.679 | 0.457 | 788.657 | 0 | 11.969 | 0.471 | 25.411 | 0 | 8813.50 | 0.458 | 155.631 | 0 |
| **ERS** | **Covariables x dtr** | w-MAT | 8.69 | 0.48 | 18.097 | 0 | 4.585 | 8812.77 | 0.679 | 0.457 | 788.657 | 0 | 8.808 | 0.472 | 18.643 | 0 | 8813.50 | 0.458 | 155.631 | 0 |
| **ERS** | **Covariables x dtr** | mat.weighted.mean:dtr.max | 2.218 | 0.324 | 6.854 | 0 | 2.879 | 8812.77 | 0.679 | 0.457 | 788.657 | 0 | 2.214 | 0.324 | 6.84 | 0 | 8813.50 | 0.458 | 155.631 | 0 |
| **ERS** | **Covariables x dtr** | log(Max. Size) | 0.668 | 0.229 | 2.921 | 0.004 | 1.079 | 8812.77 | 0.679 | 0.457 | 788.657 | 0 | 0.678 | 0.219 | 3.103 | 0.002 | 8813.50 | 0.458 | 155.631 | 0 |
| **ERS** | **Covariables x dtr** | w-TAP | 0.314 | 0.274 | 1.147 | 0.252 | 1.637 | 8812.77 | 0.679 | 0.457 | 788.657 | 0 | 0.269 | 0.27 | 0.999 | 0.318 | 8813.50 | 0.458 | 155.631 | 0 |
| **ERS** | **Covariables x dtr** | tap.weighted.mean:dtr.max | 0.89 | 0.405 | 2.195 | 0.028 | 3.134 | 8812.77 | 0.679 | 0.457 | 788.657 | 0 | 0.836 | 0.405 | 2.066 | 0.039 | 8813.50 | 0.458 | 155.631 | 0 |
| **ERS** | **Covariables x ATR** | Intercept | 26.301 | 0.532 | 49.416 | 0 | NA | 9107.56 | 0.567 | 0.318 | 493.869 | 0 | 26.092 | 0.321 | 81.267 | 0 | 9108.78 | 0.318 | 86.135 | 0 |
| **ERS** | **Covariables x ATR** | log(Max. Abund.) | 2.469 | 0.263 | 9.393 | 0 | 1.212 | 9107.56 | 0.567 | 0.318 | 493.869 | 0 | 2.459 | 0.26 | 9.456 | 0 | 9108.78 | 0.318 | 86.135 | 0 |
| **ERS** | **Covariables x ATR** | Max. ATR | 9.125 | 0.539 | 16.941 | 0 | 4.797 | 9107.56 | 0.567 | 0.318 | 493.869 | 0 | 9.138 | 0.537 | 17.01 | 0 | 9108.78 | 0.318 | 86.135 | 0 |
| **ERS** | **Covariables x ATR** | w-MAT | 2.133 | 0.413 | 5.16 | 0 | 2.625 | 9107.56 | 0.567 | 0.318 | 493.869 | 0 | 2.39 | 0.401 | 5.956 | 0 | 9108.78 | 0.318 | 86.135 | 0 |
| **ERS** | **Covariables x ATR** | mat.weighted.mean:ATR.max | 3.435 | 0.442 | 7.777 | 0 | 5.825 | 9107.56 | 0.567 | 0.318 | 493.869 | 0 | 3.449 | 0.443 | 7.793 | 0 | 9108.78 | 0.318 | 86.135 | 0 |
| **ERS** | **Covariables x ATR** | log(Max. Size) | 1.282 | 0.256 | 5.002 | 0 | 1.06 | 9107.56 | 0.567 | 0.318 | 493.869 | 0 | 1.303 | 0.243 | 5.366 | 0 | 9108.78 | 0.318 | 86.135 | 0 |
| **ERS** | **Covariables x ATR** | w-TAP | 2.712 | 0.357 | 7.593 | 0 | 2.201 | 9107.56 | 0.567 | 0.318 | 493.869 | 0 | 2.593 | 0.352 | 7.368 | 0 | 9108.78 | 0.318 | 86.135 | 0 |
| **ERS** | **Covariables x ATR** | tap.weighted.mean:ATR.max | 0.89 | 0.414 | 2.147 | 0.032 | 3.878 | 9107.56 | 0.567 | 0.318 | 493.869 | 0 | 0.831 | 0.415 | 2.004 | 0.045 | 9108.78 | 0.318 | 86.135 | 0 |
| **SES** | **Elev.** | Intercept | 0.131 | 0.081 | 1.614 | 0.107 | NA | 3550.91 | 0.122 | 0.007 | 10.298 | 0.001 | -0.004 | 0.027 | -0.145 | 0.884 | 3567.80 | 0.014 | 19.37 | 0 |
| **SES** | **Elev.** | Elev. Midpoint | -0.101 | 0.031 | -3.246 | 0.001 | NA | 3550.91 | 0.122 | 0.007 | 10.298 | 0.001 | -0.12 | 0.027 | -4.401 | 0 | 3567.80 | 0.014 | 19.37 | 0 |
| **SES** | **Max. Size** | Intercept | 0.159 | 0.087 | 1.819 | 0.069 | NA | 3508.47 | 0.207 | 0.041 | 52.739 | 0 | -0.004 | 0.027 | -0.148 | 0.883 | 3531.14 | 0.042 | 57.062 | 0 |
| **SES** | **Max. Size** | log(Max. Size) | 0.212 | 0.029 | 7.336 | 0 | NA | 3508.47 | 0.207 | 0.041 | 52.739 | 0 | 0.203 | 0.027 | 7.554 | 0 | 3531.14 | 0.042 | 57.062 | 0 |
| **SES** | **Max. Abund.** | Intercept | 0.124 | 0.086 | 1.445 | 0.149 | NA | 3418.66 | 0.316 | 0.106 | 142.548 | 0 | -0.004 | 0.026 | -0.152 | 0.879 | 3452.75 | 0.099 | 141.381 | 0 |
| **SES** | **Max. Abund.** | log(Max. Abund.) | 0.321 | 0.026 | 12.275 | 0 | NA | 3418.66 | 0.316 | 0.106 | 142.548 | 0 | 0.309 | 0.026 | 11.89 | 0 | 3452.75 | 0.099 | 141.381 | 0 |
| **SES** | **w-MAT** | Intercept | 0.142 | 0.084 | 1.692 | 0.091 | NA | 3559.14 | 0.079 | 0.001 | 2.069 | 0.15 | -0.004 | 0.027 | -0.145 | 0.885 | 3579.12 | 0.005 | 7.953 | 0.005 |

| | | | | | | | | | | | | | | | | | | | | |
|---|---|---|---|---|---|---|---|---|---|---|---|---|---|---|---|---|---|---|---|---|
| **SES** | **w-MAT** | w-MAT | 0.045 | 0.031 | 1.458 | 0.145 | NA | 3559.14 | 0.079 | 0.001 | 2.069 | 0.15 | 0.077 | 0.027 | 2.82 | 0.005 | 3579.12 | 0.005 | 7.953 | 0.005 |
| **SES** | **w-TAP** | Intercept | 0.152 | 0.087 | 1.745 | 0.081 | NA | 3561.06 | -0.016 | 0 | 0.143 | 0.706 | -0.004 | 0.027 | -0.144 | 0.885 | 3586.73 | -0.001 | 0.329 | 0.566 |
| **SES** | **w-TAP** | w-TAP | -0.011 | 0.028 | -0.381 | 0.703 | NA | 3561.06 | -0.016 | 0 | 0.143 | 0.706 | 0.016 | 0.027 | 0.574 | 0.566 | 3586.73 | -0.001 | 0.329 | 0.566 |
| **SES** | **Max. dtr** | Intercept | 0.202 | 0.105 | 1.912 | 0.056 | NA | 3489.78 | 0.167 | 0.057 | 71.425 | 0 | -0.004 | 0.027 | -0.146 | 0.884 | 3550.92 | 0.027 | 36.594 | 0 |
| **SES** | **Max. dtr** | Max. dtr | 0.261 | 0.03 | 8.764 | 0 | NA | 3489.78 | 0.167 | 0.057 | 71.425 | 0 | 0.164 | 0.027 | 6.049 | 0 | 3550.92 | 0.027 | 36.594 | 0 |
| **SES** | **Max. ATR** | Intercept | 0.181 | 0.102 | 1.776 | 0.076 | NA | 3472.78 | 0.206 | 0.07 | 88.43 | 0 | -0.004 | 0.027 | -0.148 | 0.883 | 3531.29 | 0.042 | 56.913 | 0 |
| **SES** | **Max. ATR** | Max. ATR | 0.277 | 0.029 | 9.701 | 0 | NA | 3472.78 | 0.206 | 0.07 | 88.43 | 0 | 0.202 | 0.027 | 7.544 | 0 | 3531.29 | 0.042 | 56.913 | 0 |
| **SES** | **Covariables** | Intercept | 0.096 | 0.075 | 1.293 | 0.196 | NA | 3367.64 | 0.391 | 0.143 | 199.566 | 0 | -0.004 | 0.025 | -0.157 | 0.876 | 3381.03 | 0.15 | 57.426 | 0 |
| **SES** | **Covariables** | log(Max. Abund.) | 0.337 | 0.028 | 12.2 | 0 | 1.168 | 3367.64 | 0.391 | 0.143 | 199.566 | 0 | 0.339 | 0.027 | 12.45 | 0 | 3381.03 | 0.15 | 57.426 | 0 |
| **SES** | **Covariables** | w-MAT | 0.154 | 0.032 | 4.768 | 0 | 1.274 | 3367.64 | 0.391 | 0.143 | 199.566 | 0 | 0.163 | 0.029 | 5.647 | 0 | 3381.03 | 0.15 | 57.426 | 0 |
| **SES** | **Covariables** | log(Max. Size) | 0.148 | 0.028 | 5.316 | 0 | 1.051 | 3367.64 | 0.391 | 0.143 | 199.566 | 0 | 0.142 | 0.026 | 5.498 | 0 | 3381.03 | 0.15 | 57.426 | 0 |
| **SES** | **Covariables** | w-TAP | 0.03 | 0.029 | 1.044 | 0.297 | 1.262 | 3367.64 | 0.391 | 0.143 | 199.566 | 0 | 0.033 | 0.029 | 1.154 | 0.249 | 3381.03 | 0.15 | 57.426 | 0 |
| **SES** | **Covariables x dtr** | Intercept | 0.146 | 0.067 | 2.169 | 0.03 | NA | 3072.17 | 0.575 | 0.322 | 501.038 | 0 | 0.03 | 0.031 | 0.986 | 0.324 | 3085.31 | 0.327 | 89.743 | 0 |
| **SES** | **Covariables x dtr** | log(Max. Abund.) | 0.251 | 0.025 | 10.031 | 0 | 1.215 | 3072.17 | 0.575 | 0.322 | 501.038 | 0 | 0.257 | 0.025 | 10.386 | 0 | 3085.31 | 0.327 | 89.743 | 0 |
| **SES** | **Covariables x dtr** | Max. dtr | 0.923 | 0.05 | 18.367 | 0 | 4.236 | 3072.17 | 0.575 | 0.322 | 501.038 | 0 | 0.924 | 0.05 | 18.387 | 0 | 3085.31 | 0.327 | 89.743 | 0 |
| **SES** | **Covariables x dtr** | w-MAT | 0.872 | 0.052 | 16.809 | 0 | 4.197 | 3072.17 | 0.575 | 0.322 | 501.038 | 0 | 0.89 | 0.05 | 17.657 | 0 | 3085.31 | 0.327 | 89.743 | 0 |
| **SES** | **Covariables x dtr** | mat.weighted.mean:dtr. max | -0.002 | 0.034 | -0.068 | 0.946 | 2.805 | 3072.17 | 0.575 | 0.322 | 501.038 | 0 | -0.01 | 0.035 | -0.276 | 0.783 | 3085.31 | 0.327 | 89.743 | 0 |
| **SES** | **Covariables x dtr** | log(Max. Size) | 0.064 | 0.025 | 2.562 | 0.011 | 1.086 | 3072.17 | 0.575 | 0.322 | 501.038 | 0 | 0.063 | 0.023 | 2.721 | 0.007 | 3085.31 | 0.327 | 89.743 | 0 |
| **SES** | **Covariables x dtr** | w-TAP | 0.059 | 0.029 | 2.015 | 0.044 | 1.62 | 3072.17 | 0.575 | 0.322 | 501.038 | 0 | 0.063 | 0.029 | 2.185 | 0.029 | 3085.31 | 0.327 | 89.743 | 0 |
| **SES** | **Covariables x dtr** | tap.weighted.mean:dtr. max | 0.109 | 0.043 | 2.537 | 0.011 | 2.985 | 3072.17 | 0.575 | 0.322 | 501.038 | 0 | 0.1 | 0.043 | 2.312 | 0.021 | 3085.31 | 0.327 | 89.743 | 0 |
| **SES** | **Covariables x ATR** | Intercept | 0.264 | 0.058 | 4.527 | 0 | NA | 3081.43 | 0.574 | 0.315 | 491.774 | 0 | 0.201 | 0.031 | 6.578 | 0 | 3087.38 | 0.326 | 89.304 | 0 |
| **SES** | **Covariables x ATR** | log(Max. Abund.) | 0.254 | 0.025 | 10.153 | 0 | 1.213 | 3081.43 | 0.574 | 0.315 | 491.774 | 0 | 0.255 | 0.025 | 10.317 | 0 | 3087.38 | 0.326 | 89.304 | 0 |
| **SES** | **Covariables x ATR** | Max. ATR | 0.861 | 0.051 | 16.818 | 0 | 4.684 | 3081.43 | 0.574 | 0.315 | 491.774 | 0 | 0.871 | 0.051 | 17.032 | 0 | 3087.38 | 0.326 | 89.304 | 0 |
| **SES** | **Covariables x ATR** | w-MAT | 0.449 | 0.04 | 11.329 | 0 | 2.547 | 3081.43 | 0.574 | 0.315 | 491.774 | 0 | 0.478 | 0.038 | 12.519 | 0 | 3087.38 | 0.326 | 89.304 | 0 |
| **SES** | **Covariables x ATR** | mat.weighted.mean:ATR. max | 0.188 | 0.042 | 4.491 | 0 | 5.739 | 3081.43 | 0.574 | 0.315 | 491.774 | 0 | 0.19 | 0.042 | 4.511 | 0 | 3087.38 | 0.326 | 89.304 | 0 |
| **SES** | **Covariables x ATR** | log(Max. Size) | 0.094 | 0.025 | 3.823 | 0 | 1.063 | 3081.43 | 0.574 | 0.315 | 491.774 | 0 | 0.096 | 0.023 | 4.142 | 0 | 3087.38 | 0.326 | 89.304 | 0 |
| **SES** | **Covariables x ATR** | w-TAP | 0.262 | 0.034 | 7.705 | 0 | 2.189 | 3081.43 | 0.574 | 0.315 | 491.774 | 0 | 0.256 | 0.033 | 7.66 | 0 | 3087.38 | 0.326 | 89.304 | 0 |
| **SES** | **Covariables x ATR** | tap.weighted.mean:ATR. max | 0.118 | 0.039 | 3.011 | 0.003 | 3.815 | 3081.43 | 0.574 | 0.315 | 491.774 | 0 | 0.112 | 0.039 | 2.834 | 0.005 | 3087.38 | 0.326 | 89.304 | 0 |
